# Comparative Genomics Points to Tandem Duplications of *SAD* Gene Clusters as Drivers of Increased ω-3 Content in *S. hispanica* Seeds

**DOI:** 10.1101/2023.08.27.555029

**Authors:** Tannaz Zare, Jeff F. Paril, Emma M. Barnett, Parwinder Kaur, Rudi Appels, Berit Ebert, Ute Roessner, Alexandre Fournier-Level

## Abstract

- A high-quality chromosome-level reference genome of *S. hispanica* was assembled and analysed.
- Ancestral whole-genome duplication events have not promoted the high α-linolenic acid content in *S. hispanica* seeds
- Tandem duplication of six stearoyl-ACP desaturase genes is a plausible cause for high ω-3 content in chia seeds.

*Salvia hispanica* L. (chia) is an abundant source of ω-3 polyunsaturated fatty acids (PUFAs) that are highly beneficial to human health. The genomic basis for this accrued PUFA content in this emerging crop was investigated through the assembly and comparative analysis of a chromosome-level reference genome for *S. hispanica* (321.5 Mbp). The highly contiguous 321.5Mbp genome assembly, which covers all six chromosomes enabled the identification of 32,922 protein coding genes. Two whole-genome duplications (WGD) events were identified in the *S. hispanica* lineage. However, these WGD events could not be linked to the high α-linolenic acid (ALA, ω-3) accumulation in *S. hispanica* seeds based on phylogenomics. Instead, our analysis supports the hypothesis that evolutionary expansion through tandem duplications of specific lipid gene families, particularly the stearoyl-acyl carrier protein (ACP) desaturase (*ShSAD*) gene family, is the main driver of the abundance of ω-3 PUFAs in *S. hispanica* seeds. The insights gained from the genomic analysis of *S. hispanica* will help leveraging advanced genome editing techniques and will greatly support breeding efforts for improving ω-3 content in other oil crops.

## 1 INTRODUCTION

*Salvia hispanica* L. (chia) is an oleaginous short-day flowering plant originating from Mexico and widely cultivated throughout Latin America, Australia, and Southeast Asia (Jamboonsri et al., 2012). Member of the Lamiaceae family, which comprises nearly 7,100 species of flowering plants, *S. hispanica* belongs to the largest genus, *Salvia* spp. (sages) which regroups more than 1,000 species. *Salvia* spp. are increasingly recognised as commercially important crops due to their nutraceutical and bioactive compounds among which sterols, flavonoids, diterpenes, triterpenes and polyphenols (Cahill, 2003; Harley et al., 2004; Walker et al., 2004; Georgiev & Pavlov, 2017).

Amongst the *Salvia* spp., *S. hispanica* is the most nutritionally valuable crop due to its health-promoting properties, including high levels of dietary fibre (35%), carbohydrates (5%), protein (18-24%), lipids (31-34%), antioxidants and essential vitamins (Timilsena et al., 2016; da Silva et al., 2017; Zare et al., 2019). *S. hispanica* seeds are rich in essential fatty acids (EFAs) (81%), including α-linolenic acid (ALA, ω-3, 62%) and linoleic acid (LA, ω-6, 19%) with low ω-6:ω-3 ratio (0.3), which makes it one of the best sources of plant-based ω-3 (Oteri et al., 2023). Several studies examining the effect of ω-3 polyunsaturated fatty acids (PUFAs) supplementation on human health suggest they may help reduce several chronic diseases such as diabetes, cardiovascular and inflammatory disorders, hypertension, dyslipidemia, and kidney dysfunction (Creus et al., 2017; Meyer & De Groot, 2017; Onneken, 2018; Arredondo-Mendoza et al., 2020; Penson & Banach, 2020; El-Feky et al., 2022; Xiao et al., 2022; Zhang et al., 2022; Liu et al., 2023; Ong et al., 2023).

The biosynthesis of PUFAs in plants involves a series of complex reactions in different subcellular compartments. The *de novo* biosynthesis of 16-or 18-carbon fatty acids (FAs) takes place in plastids through the action of acetyl-CoA carboxylase (ACC) and FA synthase (FAS) (Li-Beisson et al., 2013). After the conversion/elongation of C16:0 to C18:0, the C18:0-ACP (SA, stearic acid) is desaturated to C18:1-ACP (OA, ω-9, oleic acid) in the chloroplast stroma by a soluble stearoyl-ACP desaturase (SAD) (Bates et al., 2013). The C18:1-ACP is further desaturated into C16:3 and C18:2/C18:3 by different plastidial membrane-bound FA desaturases (FAD5, FAD6, FAD7/FAD8) (Browse & Somerville, 1991). FAs are next exported to the endoplasmic reticulum (ER) for conversion into acyl-CoAs before forming phosphatidylcholines (PCs) and triglycerides (TGs) (Block & Jouhet, 2015). Once synthesised, TGs are assembled into oil bodies and exported from the ER to be stored in the seed (Banaś et al., 2013).

High-quality genomes are providing valuable information on the evolution and functional divergence of key genes involved in oil biosynthetic pathways (Wang et al., 2014; Badouin et al., 2017; Unver et al., 2017; Lin et al., 2022; Shen et al., 2022). Expansion of the type 1 lipid transfer (*LTP1*) gene family and contraction of lipid degradation genes have been linked to the high oil accumulation in sesame seeds (Wang et al., 2014). Neo-functionalization and expansion of the *SAD* gene family is thought to be responsible for the increased levels of OA in olives (Unver et al., 2017). However, the lack of sufficient genomic information for *S. hispanica* had limited the exploration of the genetic basis of ω-3 PUFAs accumulation in this plant.

Early research determined chia’s somatic chromosome number and DNA content (2n = 2x = 12, C-value = 0.93 ± 0.016 pg, genome size = ∼ 460 Mb) (Haque, 1980; Estilai et al., 1990; Maynard & Ruter, 2022). In recent years, several studies have provided multi-tissue transcriptomes for *S. hispanica* in order to identify genes involved in secondary metabolite and oil biosynthesis (Sreedhar et al., 2015; Peláez et al., 2019; Wimberley et al., 2020; Gupta et al., 2021). In addition, a set of studies functionally characterised genes encoding fatty acid desaturases (FADs) against different biotic/abiotic stresses (Xue et al., 2018) (Xue et al., 2023). These studies, together with a genome assembly for *S. hispanica* (Wang et al., 2022), provided new insights, but relatively little is known about the main drivers of high ω-3 PUFA accumulation in *S. hispanica* seeds.

Our study investigated the molecular mechanisms of oil biosynthesis in *S. hispanica* leveraging the assembly of a near-complete, high-quality chromosome-level reference genome (RefSeq: GCF_023119035.1). This enabled comparative genomic analysis to determine the occurrence of WGD events and gene family size and sequence evolution between *S. hispanica* and a subset of relevant species species. We investigated if specific biological functions overrepresented among significantly expanded gene families. In particular, our analysis seek to probe the hypothesis that duplication and nucleotides substitutions in oil biosynthesis genes support the high production of ω-3 FAs in *S. hispanica*.

## 2 MATERIALS AND METHODS

### 2.1 Plant material and genomic DNA extraction

A black-seed variety of *S. hispanica* L. was sourced from Chia Co. and Northern Australia Crop Research Alliance (NACRA; Kununurra, Western Australia). Fresh young leaves were harvested from a four-week-old individual *S. hispanica* plant, immediately frozen in liquid nitrogen and stored at −80 °C prior to the isolation of genomic DNA (gDNA). High molecular weight gDNA was isolated using a cetyltrimethylammonium bromide (CTAB) method (Murray & Thompson, 1980; Supplemental Figure S1). The isolated gDNA was treated with RNase A following the method developed by Yoshinaga & Dalin (Yoshinaga & Dalin, 2016); and purified using NucleoMag^™^ NGS magnetic beads (Macherey-Nagel, Düren, Germany) prior to DNA libraries synthesis.

### 2.2 Library construction, sequencing, and processing of the sequencing reads

For short-read sequencing, DNA libraries were synthesised from 3.9 μg of gDNA using the Illumina TruSeq DNA PCR-Free kit (Illumina, San Diego, CA, USA) and sequenced on Illumina NovaSeq 6000 platform (Illumina, San Diego, CA, USA) in 2×150bp sequencing mode. Genewiz (Suzhou, China) conducted the Illumina library synthesis and sequencing. The reads quality was assessed using FastQC v0.11.9 (Andrews, 2010). Low-quality reads with an average quality per base below Q20 calculated over 4bp sliding windows and leading bases with a quality score below Q20 were removed using Trimmomatic v0.39 (Supplemental Table S1; Bolger et al., 2014). A total of 476Gb of high-quality Illumina reads with an average length of 145bp was retained for genome assembly.

For long-read sequencing, DNA libraries were prepared using the SQK-LSK109 ligation sequencing kit (Oxford Nanopore Technologies, Oxford, UK) and sequenced on a MinION Mk1B portable device with FLO-MIN106D flowcell. The long-read sequencing was run for 48hrs at 180mV using the MinKNOW software v.2.0. Basecalling of long sequencing reads was performed with Guppy v5.0.11+2b6dbffa5 using the basecalling template_r9.4.1_450bps_hac.jsn (Oxford Nanopore community, https://community.nanoporetech.com). Long reads were error-corrected with fmlrc2 v0.1.5 (Wang et al., 2018) resulting in 9Gb of high-quality resds with average length 2,825bp.

For Hi-C sequencing, nuclei were isolated from young leaves of an individual *S. hispanica* plant, and *in situ* Hi-C library synthesis was performed by DNA Zoo at the University of Western Australia (Perth, Australia) as described in Rao et al. (2014; Supplemental Figure S2). The sequencing of the Hi-C libraries (∼300bp insert size) was carried out on an Illumina NovaSeq 6000 platform (Illumina, San Diego, CA, USA) in the 2×150bp mode by Genewiz (Suzhou, China).

### 2.3 Estimation of the genome size and genomic heterozygosity

The genome size of *S. hispanica* was estimated through k-mer frequency analysis of the sequencing reads. The k-mer distributions for size ranging from 15-mer to 21-mer were computed using Jellyfish v2.3.0 (Marçais & Kingsford, 2011) and the genome size, level of the heterozygosity, and abundance of genomic repeats were estimated using GenomeScope v1.0.0 (Vurture et al., 2017).

### 2.4 *De novo* genome assembly and scaffolding

A meta-assembly approach was conducted using a hybrid combination of long and short reads. The hybrid assembly consisted in combining the contigs assembled from short-reads and error-corrected long reads using Platanus-allee with default parameters v2.2.2 (Kajitani et al., 2019). The error-corrected long reads were also used to generate a long-read only assembly using Wtdbg2 v2.5 (Ruan & Li, 2020) with default parameters. Long-read only assembly and the consensus scaffolds from the hybrid assembly were integrated into a non-redundant meta-assembly using QuickMerge v0.3 (Chakraborty et al., 2016) with the parameters “-hco 5.0 -c 1.5 −l 1000 -ml 8000 -t 16”. Iterative polishing was performed using Racon v1.4.22 (Vaser et al., 2017) with Illumina short and corrected long reads sequencing data. Ambiguous regions (N’s) and gaps within contigs were filled using Cobbler v0.6.1 (Warren, 2016). The gap-free contigs were then re-merged using RAILS v1.5.1 (Warren, 2016), and duplicated regions (haplotigs) were purged using purge_dups v1.2.5 (Guan et al., 2020) to remove misassembled or redundant contigs from the final set of haplotigs retained in the assembly. The final contig assembly was obtained after one round of Illumina short-read polishing and two rounds of corrected long-read polishing with Racon v1.4.22 (Vaser et al., 2017). The final contig assembly was subsequently scaffolded with Hi-C reads using the Juicer pipeline (Durand et al., 2016). The Hi-C-based contact map was constructed using 3D-DNA v180419 (Dudchenko et al., 2017) and manually curated using the JuiceBox v1.11.08 (Durand et al., 2016).

### 2.5 Assessment of the assembly completeness

Short and long reads were mapped to the assembled genome using bwa-mem v0.7.17 (Li & Durbin, 2009). Genome completeness was evaluated using Benchmarking Universal Single-Copy Orthologs (BUSCO) v5.2.2 (Simão et al., 2015) with different databases: “eukaryota_odb10”, “eudicots_odb10”, “viridiplantae_odb10”, and “embryophyta_odb10”. Published transcriptomes from different tissues of *S. hispanica* (Sreedhar et al., 2015; Peláez et al., 2019; Wimberley et al., 2020; Gupta et al., 2021; Klein et al., 2021) were mapped to the assembled genome using blastn v2.10.1 (Camacho et al., 2009) to further validate the completeness of the assembly.

### 2.6 Genome annotation

The genome of *S. hispanica* was annotated using the National Center for Biotechnology Information (NCBI) Eukaryotic Genome Annotation Pipeline (Pruitt et al., 2007, 2014; O’Leary et al., 2016). Transposable elements (TEs) were identified by constructing a *de novo* library of repetitive sequences based on the assembled genome using the RepeatModeler v2.0.3 (Flynn et al., 2020). The generated library was then used to classify the TEs and tandem genomic repeats and to mask the low complexity sequences within the genome using RepeatMasker v4.1.2 (Tarailo-Graovac & Chen, 2009). Annotation features over the genome assembly were visualised as a circos plot generated by pyCircos v0.3.0 (https://github.com/ponnhide/pyCircos) and Matplotlib package v3.5.1 (Hunter, 2007).

### 2.7 Comparative genomics analysis

Comparative analysis of the *S. hispanica* genome was performed against that of *Salvia splendens* (scarlet sage) as a closely related species, *Sesamum indicum* (sesame) and *Erythranthe guttata* (monkey flower) as representatives of the Lamiales order, *Solanum lycopercicum* (tomato) as a relatively close species with a high quality genome; and *Arabidopsis thaliana* (thale cress) and *Vitis vinifera* (wine grape) as outgroups. The reference genome sequences and annotations for these species were retrieved from NCBI (Sayers et al., 2021). The comparative genome analysis in this study was conducted using the compare_genomes analysis pipeline (Paril et al., 2022; Paril et al., 2023).

Protein sequences from *S. hispanica* and the species compared were used to define gene families or orthogroups as clusters of homologous genes using OrthoFinder v2.5.4 (Emms & Kelly, 2019). The hmmsearch function from HMMER v3.3.2 (Mistry et al., 2013) was used to search for gene families that corresponded to the orthogroups identified in the Protein Analysis Through Evolutionary Relationships (PANTHER, http://pantherdb.org) gene family database using PantherHMM v16.0 (Mi et al., 2021). Evolutionary relationships between gene families and contraction and expansion of gene families across species were tested using CAFE5 v5.0 (Mendes et al., 2020). Gene enrichment among the expanded or contracted gene families was analysed using the Gene Ontology (GO) enrichment analysis tool (Ashburner et al., 2000) from the Universal Protein Resource (UniProt) database (Bateman et al., 2020).

The orthogroups containing only single-copy orthologs across every species (one-to-one orthologs) were aligned with MACSE v2.06 (Ranwez et al., 2011) and used to construct a phylogenetic tree through maximum likelihood as implemented in IQ-TREE v2.0.7 (Minh et al., 2020). IQ-TREE software was also used to estimate site-specific evolutionary rates and divergence times between species through an empirical Bayes approach. Divergence times between *A. thaliana* and *V. vinifera* (115 million years, MYA) and *S. indicum* and *S. lycopersicum* (82 MYA) were inferred from the TimeTree of Life database (http://timetree.org; Kumar et al., 2017). The divergence time between *S. splendens* and *S. hispanica* (9.6 MYA) was retrieved from Wang et al. (2022). The rates of nucleotide substitution amongst pairs of paralogs/orthologs were measured using the third codon transversion rates at four-fold degenerative (synonymous) sites (4DTv) to estimate the likelihood of WGD events. The 4DTv values were calculated based on the alignment of each pair of CDSs within orthogroups across the selected species using MACSE v2.06 (Ranwez et al., 2011).

### 2.8 Analysis of oil biosynthesis genes

The evolution of key lipid biosynthesis pathway enzymes was compared between *S. hispanica* and *S. splendens*, *S. indicum*, *E. guttata*, *S. lycopercicum*, *V. vinifera* and *A. thaliana*. We focused on 35 well-characterized genes involved in lipid and FAs biosynthesis pathways retrieved from NCBI and UniProt (Supplemental Table S2). The protein sequences encoded by these lipid pathway genes were queried against the protein sequences of genes annotated in our focal species using blastp (E-value ≤1e^-4^) to identify the orthogroups encoding for a specific gene activity. Contraction and expansion of these gene families and rates of nucleotide substitution were tested as described in the previous section.

Gene duplication events, including WGD, tandem duplication, proximal duplication, transposed duplication, and dispersed duplication, were identified using the DupGen_finder pipeline (Qiao et al., 2019). Protein sequences were first aligned using blastp with e-value < 1e^-5^, and the different modes of gene duplications between homologous gene pairs determined using the DupGen_finder.pl function from MCScanX (Qiao et al., 2019) with the following parameters were used: match_score: 50, match_size: 5, gap_penalty: −1, overlap_window: 5, e_value: 1e^-5^, max gaps: 25. The chromosome ideogram plot and homologous synteny blocks were generated using the R\RIdeogram package (Hao et al., 2020).

The ratio between the number of nonsynonymous substitutions per nonsynonymous site (Ka) and the number of synonymous substitutions per synonymous site (Ks) in a pairwise alignment of two orthologous sequences was used to measure evolutionary differences between sequences. The KaKs_Calculator2.0 v2.0 (Wang et al., 2010) was used to calculate Ka/Ks ratio over 15bp sliding windows of the CDSs of paralogs associated with lipid metabolism in the *S. hispanica* genome and homologous genes in other species.

## 3 RESULTS

### 3.1 Chromosome-scale reference genome assembly and annotation of *S. hispanica*

The genome assembled for *S. hispanica* (RefSeq: GCF_023119035.1) consisted in 5,304 contigs spanning 1,556 scaffolds. The assembly covered ∼321Mb (N50=53Mb; L50=3; largest scaffold=57Mb) with a GC content of 36% (Table 1). Hi-C reads analysis identified ∼173 million contacts (Supplemental Table S3), of which ∼127 million and ∼46 million were inter- and intra-chromosomal contacts used for genome super-scaffolding, respectively. The size of *S. hispanica* pseudo-chromosomes obtained through Hi-C scaffolding ranged from 40Mb to 58Mb with spanned gaps of 491bp to 741bp (Supplemental Table S4); The L90=6 matched the chromosome number determined through flow cytometry (Figure 1 and Supplemental Figure S3; Maynard & Ruter, 2022). The best k-mer distribution model was obtained for k=19 and supported diploidy with 0.24% heterozygosity and 5.28% duplicated regions (Supplemental Figure S4). Analysis of k-mer frequencies estimated a haploid genome size of 466Mbp, consistent with the size of 460Mbp reported by Maynard and Ruter (2022).

**Figure 1.**
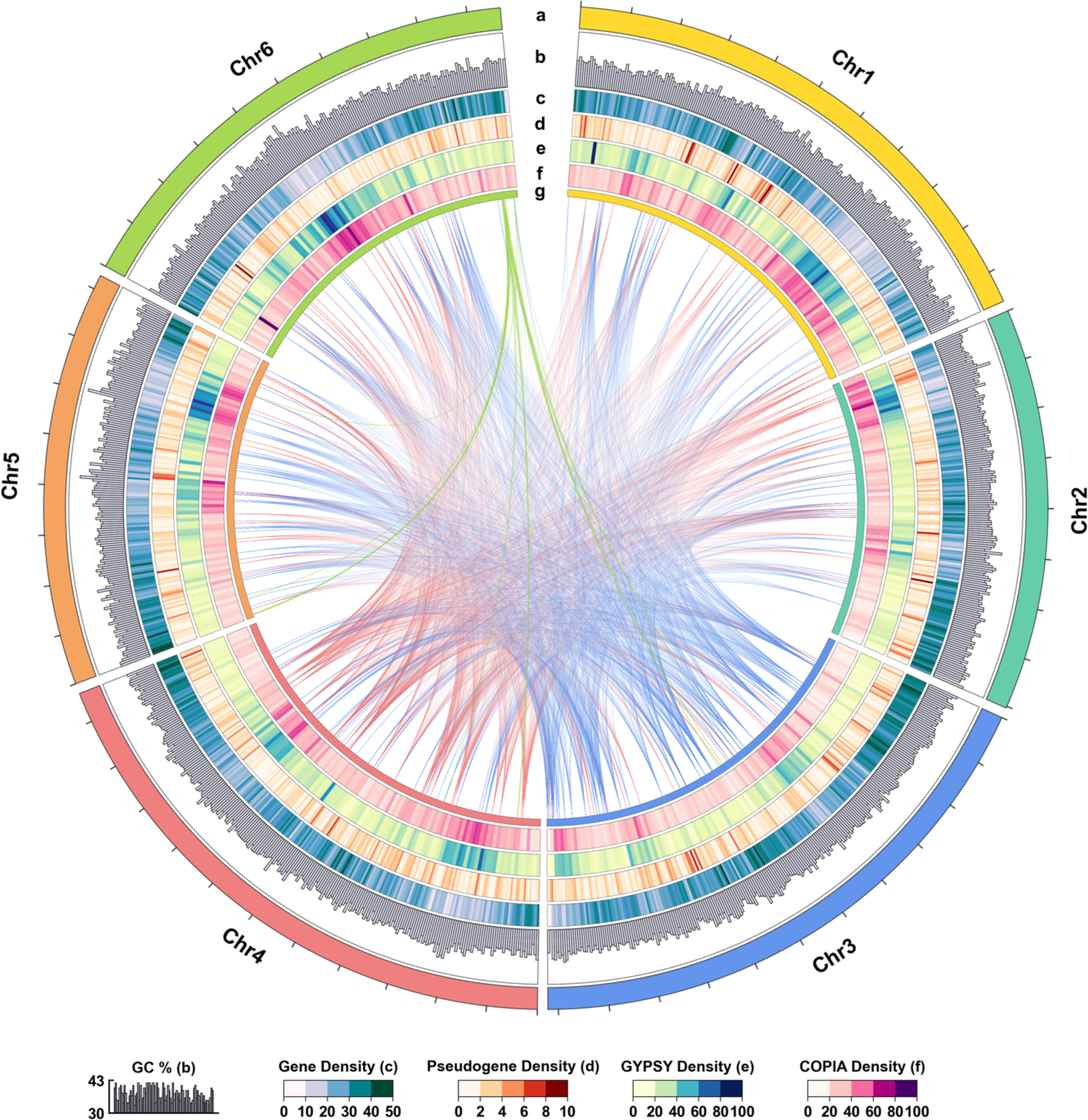
Genomic features of the *S. hispanica* reference genome assembly. (a) Chromosome layer showing the length of each chromosome with ticks indicating 5Mbp intervals. (b) Distribution of the GC content at a window size of 250kbp over the entire genome as a bar plot of percentage values with a lower bound of 30% and upper bound of 43%. (c) Distribution of protein coding gene density over 250kbp windows (values normalized between 0 and 50 across chromosomes). (d) Distribution of pseudogene density over 250kbp windows (values normalized between 0 and 10 for all chromosomes). (e) and (f) Distribution of Gypsy and Copia LTR density, respectively, over 500kbp windows (values normalized between 0 and 100 across chromosomes). (g) The chord plot shows the synteny relationships for the top five orthogroups (paralogs) across the genome. The color of the internal chords is that of the chromosome containing the highest number of paralogs within each orthogroup.

**Table 1.** Statistics for the chromosome-level reference genome assembly of *S. hispanica*.

| Assembly Features | *S. hispanica* Genome Assembly |
| --- | --- |
| Total assembly length (bp) | 321,469,233 |
| Contigs ≥ 1k bp | 5,301 |
| Contigs ≥ 10 kbp | 4,307 |
| Contigs ≥ 25 kbp | 4,016 |
| Contigs ≥ 50 kbp | 3,858 |
| Largest contig length (kbp) | 894 |
| Scaffolds ≥ 1 kbp | 1,553 |
| Scaffolds ≥ 10 kbp | 572 |
| Scaffolds ≥ 25 kbp | 288 |
| Scaffolds ≥ 50 kbp | 130 |
| Largest Scaffold length (kbp) | 57,985 |
| % Main genome in scaffolds > 50 kbp | 95.66 |
| GC (%) | 36 |
| N50 | 53,190,533 |
| L50 | 3 |
| N90 | 40,175,719 |
| L90 | 6 |
| Mismatches (N's) | 613,150 |
| Gap (%) | 0.19 |

The completeness of the assembly assessed against a different set of lineage-specific core eukaryotic genes (Eukaryota n=255, Eudicots n=2117, Embryophyte n=1538, and Viridiplantae n=410), resulted in the retrieval of 98.4%, 93.6%, 95.3%, and 96.5% of complete single copy gene models, respectively (Supplemental Figure S5). The average BUSCO score was relatively high (> 95%) across all lineage sets. Around 94% of the previously published *S. hispanica* transcripts (Sreedhar et al., 2015; Peláez et al., 2019; Wimberley et al., 2020; Gupta et al., 2021; Klein et al., 2021) mapped to the assembled genome (Supplemental Figure S6). The high BUSCO score and the high mapping rate of transcripts indicated that the genome assembly contained nearly all the *S. hispanica* genes.

Additionally, 97.56% of the short reads re-mapped against the assembled genome indicating the high quality of the *S. hispanica* reference genome. However, 0.6% of the reads did not have their paired read mapped to the genome and 5.9% of paired reads were mapped to a different chromosome. This potentially highlights repetitive sequences in the assembled genome and closely matches the estimated genome duplication rate of 5.28% determined by GenomeScope.

A total of 46,508 CDSs were annotated, encompassing 209,379 exons and 166,729 introns across all transcripts including mRNAs, misc_RNAs, and ncRNAs of class lncRNA. The repeat-masked assembly contained 39,616 genes, corresponding to 32,922 protein-coding genes, 4,071 non-coding genes and 2,623 pseudogenes (Supplemental Table S5). The total number of annotated transcripts (54,009) included 46,423 messenger RNAs (mRNAs), 3,758 long non-coding RNAs (lncRNAs), 739 transfer RNAs (tRNAs), 436 small nucleolar RNAs (snoRNAs), 233 small nuclear RNAs (ncRNAs), 49 ribosomal RNAs (rRNAs), and 2,381 miscellaneous RNAs (misc_RNAs; Table 2).

**Table 2.** Count and length of the annotated genomic features in the *S. hispanica* genome (excluding pseudogenes).

| Feature | Count | Mean<br>length<br>(bp) | Median<br>length<br>(bp) | Min<br>length<br>(bp) | Max<br>length<br>(bp) |
| --- | --- | --- | --- | --- | --- |
| Genes | 36,993 | 2,901 | 2,291 | 62 | 163,900 |
| All transcripts | 54,009 | 1,671 | 1,452 | 62 | 16,722 |
| mRNA | 46,423 | 1,753 | 1,515 | 165 | 16,722 |
| Misc_RNA | 2,381 | 2,122 | 1,859 | 167 | 13,008 |
| tRNA | 739 | 74 | 73 | 71 | 93 |
| lncRNA | 3,758 | 979 | 729 | 78 | 5,754 |
| snoRNA | 436 | 106 | 103 | 62 | 229 |
| snRNA | 223 | 138 | 120 | 98 | 197 |
| rRNA | 49 | 384 | 119 | 103 | 3,191 |
| Single exon transcripts | 5,396 | 1,149 | 957 | 233 | 6,688 |
| CDSs | 46,508 | 1,379 | 1,155 | 90 | 16,188 |
| Exons | 209,379 | 302 | 162 | 2 | 7,672 |
| Exons in coding transcripts | 197,070 | 302 | 161 | 2 | 7,062 |
| Exons in non-coding transcripts | 19,006 | 274 | 153 | 2 | 7,672 |
| Introns | 166,729 | 355 | 142 | 30 | 99,611 |
| Introns in coding transcripts | 158,426 | 343 | 139 | 30 | 99,611 |
| Introns in non-coding transcripts | 14,710 | 437 | 196 | 32 | 63,506 |

The genome of *S. hispanica* contained 44.14% of interspersed repeats, 0.80% and 0.19% of which were simple and low complexity repeats, respectively (Supplemental Table S6 and Supplemental Figure S7). Long terminal repeats (LTRs) represented 11.05% of the genome, with Copia (5.60%) and Gypsy (5.45%) being the most abundant (Figure 1h), when DNA transposons only represented 3.69% of the genome (Supplemental Table S6 and Supplemental Figure S7).

3.2 Gene family evolution in the *S. hispanica* genome

The CDSs from the *S. hispanica* genome annotation were compared to those of *S. splendens*, *S. indicum*, *E. guttata*, *S. lycopercicum*, *V. vinifera* and *A. thaliana* (Figure 2). The phylogeny inferred from 134 single-copy orthologs (SCOs) supported previously described evolutionary relationships among species. The SCOs alignment placed the *S. splendens* and *S. hispanica* with maximum nodal support, confirming their close ancestral relationship (Figure 2a). Their most recent common ancestor was dated to 9.6 million years ago (MYA) which supported previous report by Wang et al. (2022). The oilseed crops *S. indicum* and *S. hispanica* were estimated to have diverged approximately 58-59 MYA, similar to the estimated divergence time between *E. guttata* and *S. hispanica* (Figure 2a).

**Figure 2.**
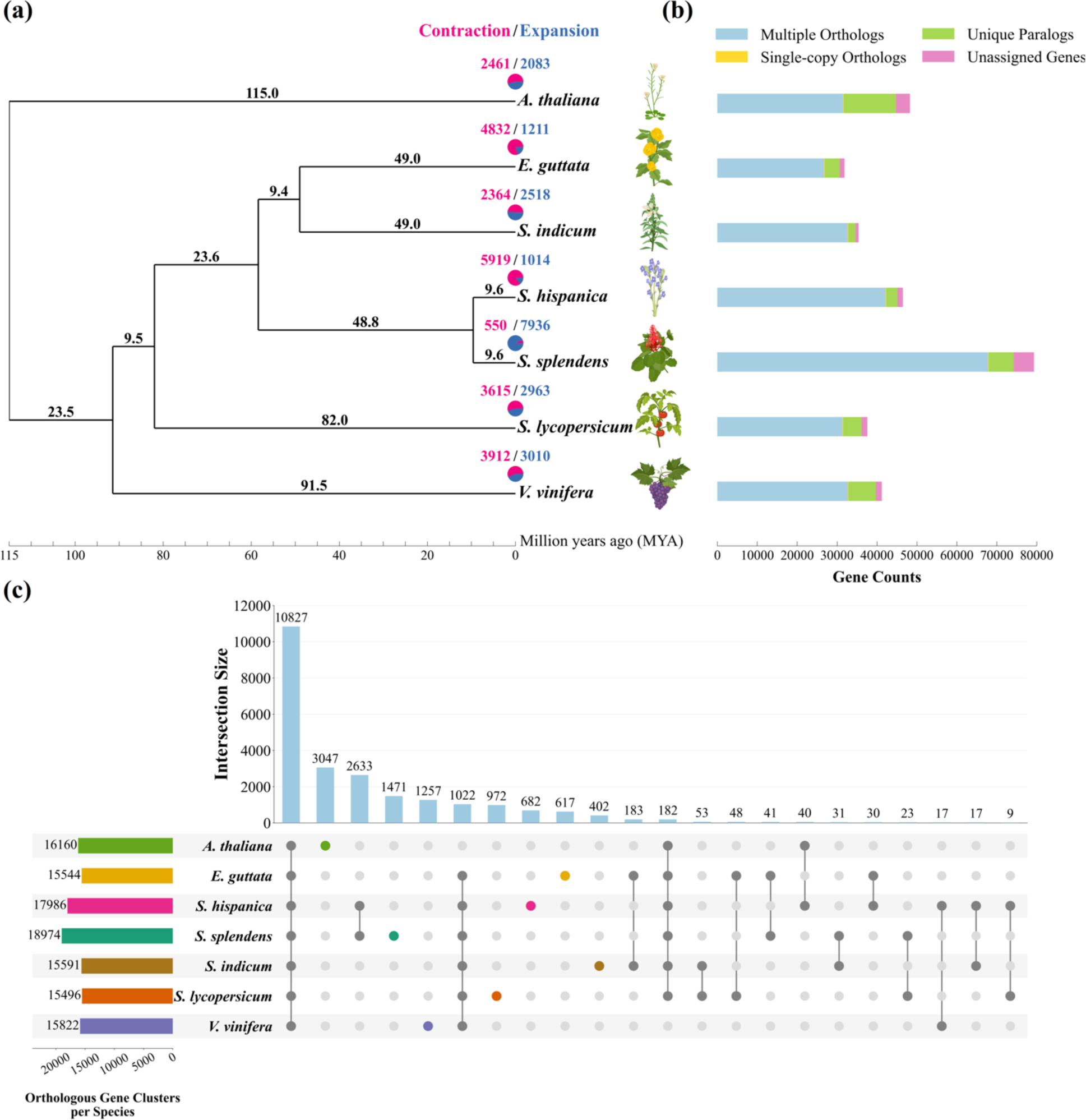
Evolution of *S. hispanica* and distribution of orthologous gene families across species. (a) Phylogenetic tree inferred from single-copy orthologs among selected species. Numbers on branches show divergence time in MYA. The pie charts at the terminal branches show the contraction (pink) and expansion (dark blue) of gene families for each species. (b) Distribution of multiple orthologs, single copy orthologs, unique paralogs and genes not associated with orthologs per species from orthogroup clustering by OrthoFinder. (c) The UpSet plot of the interactions between unique and shared orthologous gene clusters identified by OrthoFinder. The horizontal bar plot on the left shows the total number of orthogroups assigned to each species. The dark dots connected by solid lines show the species that include in each cluster where the number of orthogroups within that cluster is indicated by the vertical bars on the top. Colored dots on the cluster map represent orthogroups unique to a species. Plant images are created with BioRender.com.

From the 320,180 genes found across all seven species, 305,234 genes (95.3%) were assigned to one or more of the 27,963 orthogroups. In *S. hispanica*, a total of 45,108 genes were assigned to orthogroups, 134 being SCOs, and 2,814 unique paralogs form 682 orthogroups, leaving 1,400 unassigned genes (Figure 2b). *S. splendens* contained the highest average number of paralogs within orthogroup (1.85), showing the highest genetic redundancy, followed by *A. thaliana* (1.12), *S. hispanica* (1.08), *V. vinifera* (0.96), *S. lycopersicum* (0.88), *S. indicum* (0.83), and *E. guttata* (0.74). The highest number of unique orthogroups was observed for *A. thaliana* (3047; 13074 paralogs), followed by *S. splendens* (1471; 6,236 paralogs), *V. vinifera* (1257; 6,892 paralogs), *S. lycopersicum* (972; 4,601 paralogs), *S. hispanica* (682; 2814 paralogs), *E. guttata* (617; 3,738 paralogs), and *S. indicum* (402; 1,790 paralogs; Figure 2b). The number of genes not assigned to any orthogroup varied across species, with the highest value for *S. splendens* (5,081) followed by *A. thaliana* (3,503), *S. lycopersicum* (1,489), *V. vinifera* (1,452), *S. hispanica* (1400), *E. guttata* (1,216), and *S. indicum* (805; Figure 2b).

In total, 10,827 orthogroups were shared among all species, with 682 orthogroups being unique to *S. hispanica* (Figure 2c). The two closely related *Salvia* species (i.e., *S. splendens* and *S. hispanica*) contained the largest number of orthogroups (18,974 and 17,986, respectively), which is 15-22% higher than that observed for other species. The high number of unique orthogroups in *A. thaliana* (3,047) reflected the distant evolutionary relationships with the other species analysed and its relevance as an outgroup (Figure 2c).

We next explored gene family expansion and contraction in *S. hispanica* compared to other selected species. *A. thaliana*, *S. indicum*, *S. lycopersicum* and *V. vinifera* showed a relatively even number of expanded *vs*. contracted gene families (Figure 2a). *E. guttata* and *S. hispanica*, on the other hand, show a much higher number of contracted gene families, while *S. splendens* was the only species with significant number of expanded gene families (7936). Interestingly, closely related *S. hispanica* exhibited the opposite pattern with a significant excess of contracted gene families (5,919).

Among the gene families expanded in *S. hispanica.,* a significant enrichment was found for maintenance of plant homeostasis, response to stress and activation of defence mechanisms (Supplemental Table S7). The top 10 gene families most unique to *S. hispanica* were highly enriched for specific biological processes: xenobiotic detoxification by transmembrane export in plasma membrane (GO:1990961; P<5.70E^-10^), xenobiotic export from cell (GO:0046618; P <5.70E^-10^), xenobiotic transport (GO:0042908; P <9.63E^-11^), plant-type primary cell wall biogenesis (GO:0009833; P<3.22E^-04^), galactose metabolic process (GO:0006012; P<9.83E^-04^), peptidyl-threonine dephosphorylation (GO:0035970; P<4.17E^-08^), toxin catabolic process (GO:0009407; P<8.70E^-07^), nucleotide-sugar transmembrane transport (GO:0015780; P<2.30E^-02^); S-glycoside catabolic process (GO:0016145; P<2.14E^-04^), and glucosinolate catabolic process (GO:0019762; P<2.14E^-04^; Supplemental Figure S8). The 20 most enriched molecular function and cellular component ontologies in the *S. hispanica* genome are presented in Supplemental Table S8 and S9, respectively.

### 3.3 Whole genome duplications and speciation events

The occurrence of WGD events in species studied was determined based on the distribution of the 4DTv among multi-copy paralogs (Supplemental Figure S9). The 4DTv distribution for *S. hispanica* (Supplemental Figure S9h) showed a high density at 0.1 (relative time to the most recent common ancestor) and at 0.3. The first peak corresponded to a relatively recent WGD event in *S. hispanica* that is only shared with closely related species *S. splendens*. However, this peak in the 4DTv distribution of *S. splendens* is masked by a very recent WGD event at 0.03 (Supplemental Figure S9h). Both *S. indicum* and *S. lycopersicum* showed similar peaks at around 0.2 and 0.4 suggesting a more recent WGD; these peaks are less apparent for *E. guttata*, *A. thaliana* and *V. vinifera* (Supplemental Figure S9h).

The pairwise comparison of the 4DTv distribution in the two *Salvia* species indicated that the speciation event between them might have occurred quite recently (9.6 MYA as shown in Figure 2a), after a common WGD event shared across all *Salvia* species (Figure 3) and before the recent WGD event private to *S. splendens*. A comparison of *S. hispanica* with *S. indicum* and *E. guttata* (Figure 3) revealed that the *S. hispanica* genome has diverged from both species at the same time, supporting the estimated divergence time of 58.4 MYA (Figure 2a) and the absence of WGD private to *S. hispanica* in the *Salvia* lineage.

**Figure 3.**
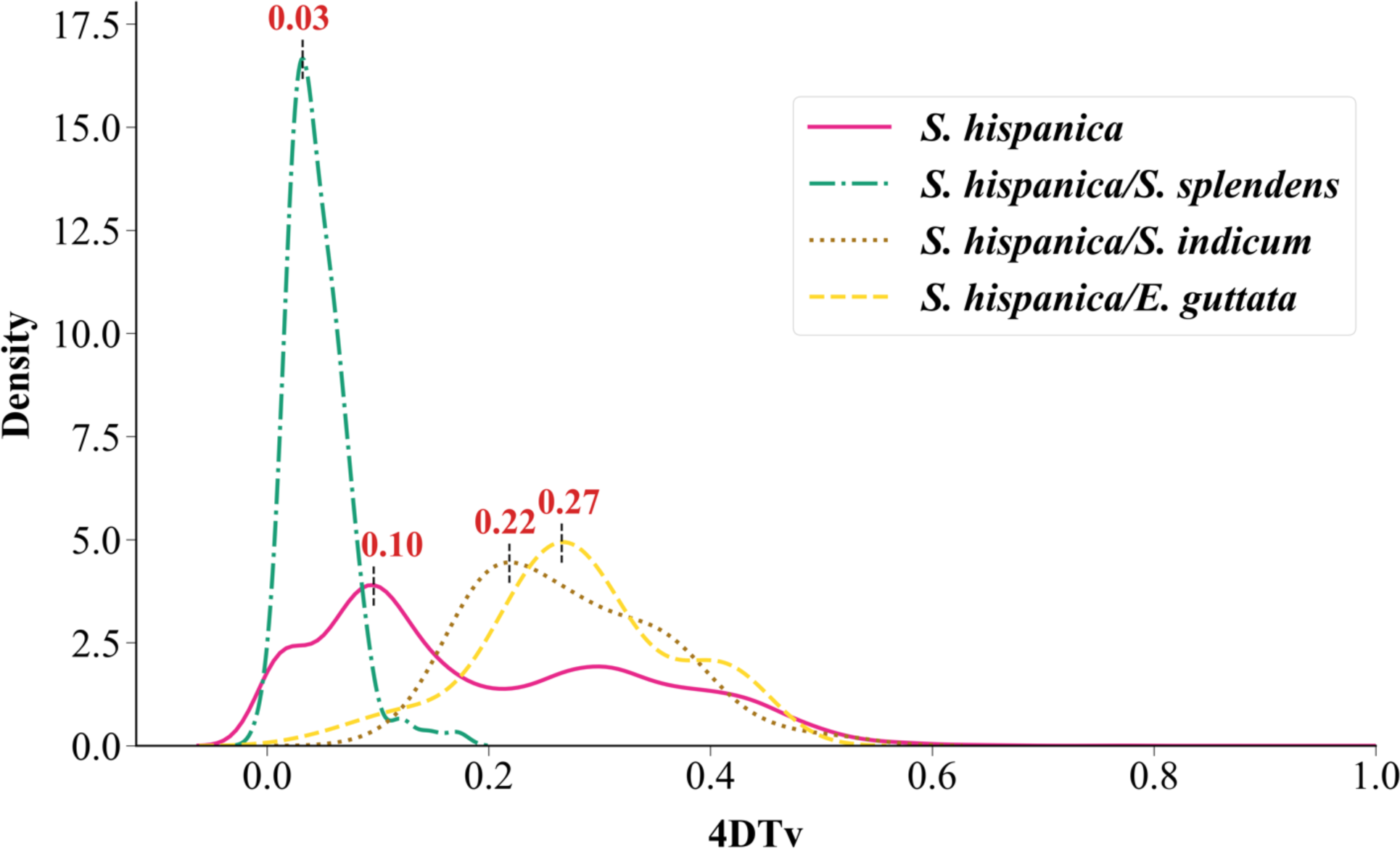
Distribution of transversion substitutions at fourfold degenerate sites (4DTv). Distribution of 4DTv for *S. hispanica* and pairwise 4DTv with *S. splendens*, *S. indicum*, and *E. guttata*. Peaks in pairwise 4DTv density indicate the relative time of divergence between species.

3.4 Analysis of oil biosynthesis genes in *S. hispanica*

We investigated gene family expansion and particularly segmental duplication as a potential hypothesis for the high production of ω-3 FAs in *S. hispanica*. Key lipid synthesis genes including *SAD* and 3-ketoacyl-acyl carrier protein reductase (*KAR*) were significantly expanded in *S. hispanica*. The increased number of *SAD* genes in *S. hispanica* cannot be explained by the WGD events: the *S. splendens* genome is twice larger than that of *S. hispanica*, having recently undergone a WGD event, but does not contain twice the number of *SAD* genes. To understand the mechanisms underlying the expansion of specific gene families in *S. hispanica*, we investigated different modes of gene duplications (i.e., whole-genome, tandem, proximal, transposed, or dispersed duplications).

In *S. hispanica*, most of the *SAD* genes are located in the telomeric region of chromosome 1 (11 out of 13 genes), and the remaining two are located on chromosomes 3 and 4 (Figure 4). Gene duplication analysis (e-value < 10^-5^) revealed that 6 of the *ShSAD* genes (*ShSAD2*: XP_047955596.1; *ShSAD3*: XP_047970931.1; *ShSAD4-a* isoform X1: XP_047970902.1; *ShSAD4-b* isoform X2: XP_047970909.1; *ShSAD5*: XP_047970920.1; *ShSAD6*: XP_047955488.1; and *ShSAD7*: XP_047954777.1) form a tandem array located in the telomeric region of chromosome 1 (Figure 4a). Interestingly, the genes in this tandem array (excluding *ShSAD7*) were specific to *S. hispanica*, absent in other species studied. One gene upstream of this tandem array, the *ShSAD1* gene (XP_047969270.1) was unique to *S. hispanica,* resulting from a duplication of *ShSAD13* (XP_047982897.1) located on chromosome 4.

**Figure 4.**
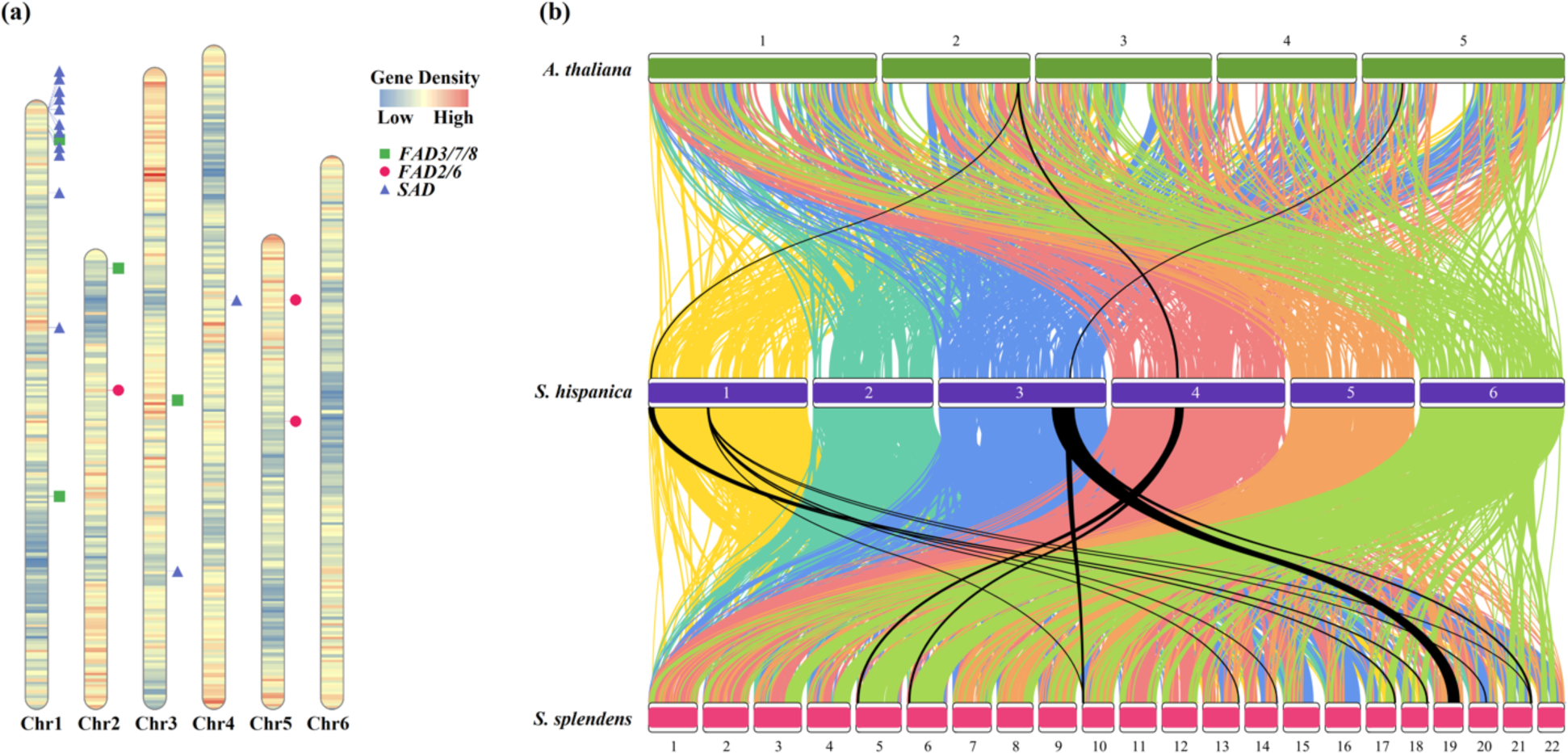
Chromosome ideogram and synteny analysis of *S. hispanica* genome. (a) Ideogram showing the gene density distribution and position of key FA synthesis genes on *S. hispanica* chromosomes. The tandem array of *ShSAD* genes is located in the telomeric region of chromosome 1. (b) Synteny analysis of *S. hispanica* with *A. thaliana* and *S. splendens* using synteny blocks from DupGen_finder. Colours represent *S. hispanica* chromosomes. Black cords represent only synteny blocks containing *ShSAD* genes.

*ShSAD13* belongs to an orthogroup shared across species that included 4 genes from *S. hispanica*. This orthogroup included *ShSAD13*, *ShSAD7*, and *ShSAD11-a* isoform X1 (XP_047955195.1), *ShSAD11-b* isoform X2 (XP_047955196.1) and *ShSAD12* (XP_047971758.1), which are dispersed duplicates located on chromosome 1 and 3, respectively. The remaining three *ShSAD* genes formed an orthogroup unique to *S. hispanica* with *ShSAD8* (XP_047966762.1) and *ShSAD9* (XP_047955361.1) which are proximal duplicates (ie. one gene apart) and *ShSAD10* (XP_047938734.1) which is a transposed duplicate of *ShSAD1*. This is different to the finding by Xue and colleagues who showed that all *ShSAD* genes are tandem duplicates (Xue et al., 2023). Instead, our analysis suggested that the *ShSAD* genes have been repeatedly duplicated in *S. hispanica,* after its divergence from *S. splendens* (Figure 4b).

The eleven *ShKAR* genes are spread across chromosomes 2 to 6, six of which directly resulted from WGD, including *ShKAR1* (XP_047958713.1), *ShKAR2* (XP_047960775.1), *ShKAR3* (XP_047964327.1), *ShKAR5* (XP_047940213.1), *ShKAR9* (XP_047939697.1), and *ShKAR10* (XP_047952057.1). *ShKAR5* is also a tandem duplicate of *ShKAR6* (XP_047944250.1) which is one gene away from the pair of tandem duplicates formed by *ShKAR7* (XP_047940355.1) and *ShKAR8* (XP_047944629.1). In addition, *ShKAR4* (XP_047937601.1) and *ShKAR11* (XP_047945899.1) are transposed duplicate and proximal duplicate pairs of *ShKAR10*, respectively.

Three characteristics were unique to the *SAD* gene family compared to the *KAR* gene family. First, the *S. hispanica* genome contained the highest number of *SAD* gene orthologs (13 copies) of all species studied here. However, the number of *KAR* family genes in *S. hispanica* was similar to that of other species. Second, *SAD* genes had the highest number of paralogs (10 genes including a private orthogroup of 3 genes) unique to *S. hispanica*, while *S. hispanica KAR* genes only showed one unique paralog. Third, *ShSAD* genes included a six-gene tandem array including 5 genes unique to *S. hispanica*, while Sh*KAR* genes included two tandem pairs with only one gene unique to *S. hispanica*.

We evaluated the evolutionary constraint on the *ShSAD* genes by analysing the Ka/Ks ratio in orthogroups containing at least two genes. Comparison of pairwise Ka/Ks ratios between CDSs of *ShSAD*s unique to *S. hispanica* and orthologs from other species revealed a set of *ShSAD* genes with functional characteristics unique to *S. hispanica*. The orthogroup containing *ShSAD8*, *ShSAD9*, and *ShSAD10* showed an excess of non-synonymous substitution, with 14 to 16% of the alignment length displaying a Ka/Ks ratio greater than 1 (Supplemental Figure 10; Fisher’s exact test: P<0.05). In contrast, Ka/Ks ratios were less than 1 for 22 to 27% of the alignments (Fisher’s exact test: P<0.05), indicative of purifying selection at other sites. However, most of the alignments length showed no substitutions, either being fully conserved (*ShSAD8*-*ShSAD9*) or not present across all species investigated (*ShSAD8*-*ShSAD10* and *ShSAD9*-*ShSAD10*). For the orthogroup containing *ShSAD7*, *ShSAD11-a/b*, *ShSAD12*, and *ShSAD13*, 68 to 90% of the alignments length showed Ka/Ks ratios lesser than 1 (Fisher’s exact test: p-value=0.05) and only 0-6% of Ka/Ks ratios greater than 1 (Fisher’s exact test: P<0.05), suggesting a strong purifying selection (Supplemental Figures 11, 12 & 13).

## 4 DISCUSSION

### 4.1 Gene family evolution in *S. hispanica*

WGD is common in angiosperms, allowing the neofunctionalization of duplicated genes and the potential adaptation to novel conditions (Hahn et al., 2005; Hughes et al., 2014; Li et al., 2020). In addition to the WGD event previously reported for *S. hispanica* (Wang et al., 2022; Li et al., 2023) shared with *S. splendens*, we identified an ancestral γ-WGD event shared with other species. *Salvia* species having undergone these two WDGs (e.g., *S. splendens*) neither show an increased expression of lipid genes nor a high oil accumulation in seeds. Therefore, the higher ω-3 production in *S. hispanica* compared to the other species studied here cannot be due to one of its WGD events.

Whole genome duplication events do not alter the dosage balance across molecular pathways, including protein modification and transcriptional regulation (Chang et al., 2022). The consistent gene expression after WGDs is due to dosage sharing and tight regulatory control mechanisms; in contrast, tandem duplications lead to the shuffling of regulatory elements (Rogers et al., 2017). Tandem duplications in *A. thaliana*, *S. lycopersicum*, and *Z. mays* were shown to impact dosage balance in protein-protein interactions and could explain that *ShSAD* genes tandem duplications increased FA synthesis in *S. hispanica* seeds. The effect of duplicated genes on dosage balance is more profound when genes encode for a limiting step of a metabolic pathway (Defoort et al., 2019), which is the case for *ShSAD* genes in FA biosynthesis.

The genome of *S. hispanica* showed the highest number of contracted gene families (5,919), when *S. splendens* showed the highest number of expanded gene families (7,936). The expansion or contraction of gene families has been associated with gene regulation (Baroncelli et al., 2016; Najafpour et al., 2020). Biological pathways are often regulated at the gene network level through regulatory hubs with key regulatory gene families expanded (Yu et al., 2017). In contrast, genes performing independent functions under purifying selection often show gene family contraction (Hess et al., 2018). Consequently, non-synonymous mutations cause immediate loss of function in single-copy genes but are neutral in expanded gene families due to functional redundancy (Force et al., 1999; Hahn et al., 2005).

The *S. hispanica* genome with predominantly contracted gene families is under purifying selection, and the selective removal of deleterious alleles potentially explains the genomic stability of key biological functions (Bray & West, 2005; Hough et al., 2013; Cvijović et al., 2018; dos Santos Maraschin et al., 2019). Intense purifying selection has been observed across plant species. In *Zea mays* (maize), highly expressed genes experience stronger purifying selection and regulatory neofunctionalization leading to unique and independent functions that have increased photosynthetic efficiency and stress tolerance (Hughes et al., 2014). Similarly, paralogs with deleterious effects might have been removed from the *S. hispanica* genome, and a structural reorganisation could have occurred within gene families involved in oil biosynthesis.

### 4.2 Tandem duplication as an essential evolutionary genomic mechanism

Tandem duplication, one of the main mechanisms of the gene family expansion (Achaz et al., 2000; Lan & Pritchard, 2016), is also supporting phenotypic plasticity (Chang et al., 2022) by mediating the adaptive response of the secondary metabolism to environmental stress (Defoort et al., 2019). Tandem duplications are lineage-specific and often affect membrane proteins and biotic and abiotic response genes (Rizzon et al., 2006; Hanada et al., 2008; Carretero-Paulet & Fares, 2012; Kondrashov, 2012; Jiang et al., 2013; Denoeud et al., 2014; Fischer et al., 2014; Picart-Picolo et al., 2020; Cai et al., 2023). In the Lamiaceae family, species-specific tandem duplicates are responsible for the biosynthesis of flavonoids in *Scutellaria baicalensis* (Xu et al., 2020a), terpenoids in the *Lavandula angustifolia* (lavender) (Li et al., 2021), and diterpenoids in *Isodon rubescens* (Sun et al., 2023).

The tandem array of six *ShSAD* genes located in the telomeric region of chromosome 1 suggests a shared regulation. Tandem duplicates are known to be co-regulated with sub-functionalization of expression at higher levels compared with segmental duplicates or WGD genes (Cannon et al., 2004; Casneuf et al., 2006). For example, tissue specific co-expression of unique tandem duplications involved in the biosynthesis of flavonoids was observed in *Carthamus tinctorius* (safflower; Wu et al., 2021). Similarly, seed specific tandem duplicated gene pairs responsible for oil biosynthesis in *S. indicum* were shown to be co-expressed (Song et al., 2021).

Tandem duplications of lipid biosynthesis genes with effects consistent with those found here in *S. hispanica* have been evidenced across different species. In *Cajanus cajan* (pigeon pea), tandem duplicates control the biosynthesis of ALA (Liu et al., 2021). In *S. indicum*, a combination of lipid transfer gene family expansion due to tandem duplication and contraction of lipid degradation genes was identified as driving the high accumulation of FAs in seeds (Wang et al., 2014). The highly conserved domains in segmental and tandem duplicated wax ester synthase (*WSD1*) and *DGAT* genes in *Gossypium hirsutum* (cotton) were related to a rate-limiting process during high unsaturated FAs accumulation in seeds (Zhao et al., 2021). The expansion of *GmFAD2* genes in *Glycine max* (soybean) (Lakhssassi et al., 2021) and *OeB3* genes in *Olea europaea* (olive) (Qu et al., 2023) were associated with tandem duplications.

### 4.3 *SAD* gene family expansion is an adaptive multi-stress response mechanism affecting FA biosynthesis

Contrary to most gene families involved in lipid synthesis in *S. hispanica*, the *ShSAD* and *ShKAR* gene families are substantially expanded. This extends previous findings from transcriptomic analysis which only showed that the *SAD* gene family was expanded (Wang et al., 2022). In plants, the *SAD* and *KAR* gene families are critical for FA biosynthesis and the initiation of FA desaturation in the chloroplast (Li-Beisson et al., 2013; González-Thuillier et al., 2021). *SAD* genes encode the only known soluble desaturase in chloroplast stroma, which is essential for ALA biosynthesis. The SAD enzyme also controls the synthesis of ACP-bound oleic acid (18:1) from stearate, resulting in the first double bond at the α-end (You et al., 2014).

In *Camellia chekiangoleosa* seeds, the expansion of the *SAD* gene family leads to the high production of unsaturated FAs, which is thought to be adaptive (Shen et al., 2022). Similarly, in *S. hispanica* seeds with high FA content, a high number of unique *SAD* genes sit in a tandem array. *SAD* genes sitting in tandem have also been reported in *Linum usitatissimum* L. (flax seeds; You et al., 2014) and *Olea europaea* var. *sylvestris* (wild olive) where the expansion of duplicated *SAD* genes has allowed neofunctionalization to support the high production of OA (Unver et al., 2017).

The increased number of tandem-duplicated *SAD* genes in the *S. hispanica* genome, leading to the abundant production of PUFAs, might represent an adaptive response to environmental stress as shown in other plants (Feng et al., 2017; Zhao et al., 2021; Chen et al., 2023). The *ShSAD2* and *ShSAD7* genes in *S. hispanica* are overexpressed in response to cold stress (Xue et al., 2023). Differential substrate specificity of tandem duplicates is believed to be the mechanism behind this adaptive response to stress also leading to enhanced secondary metabolite synthesis (Wang et al., 2015; Picart-Picolo et al., 2020; Tohge & Fernie, 2020; Xiao et al., 2020; Xu et al., 2020b; Li et al., 2021; Chang et al., 2022).

The function of *ShSAD11a* in seed oil formation was confirmed through heterologous expression studies in yeast and *A. thaliana* transgenic lines (Xue et al., 2023). The higher number of *SAD* genes in *S. hispanica* (13 genes) compared to *Perilla frutescens* (7 genes) is currently the main hypothesis for the higher accumulation of ALA in *S. hispanica* seeds (Xue et al., 2023). However, we also specifically hypothesise that the six-gene tandem array of *ShSAD* genes in the telomeric region of chromosome 1, including five species-specific, co-expressed genes might further explain the high accumulation of ω-3 FAs in *S. hispanica* seeds. For example, regulation of the very long-chain monounsaturated nervonic acid (NA, C24:1ω-9) in *Acer truncatum* (purple blow maple) is controlled by a 10-gene tandem array of 3-ketoacyl CoA synthetase (*KCS*) genes, encoding a rate-limiting enzyme defining substrate and tissue specificity during FA elongation, and highly expressed in mature seeds (Ma et al., 2020).

The *ShSAD* tandem array includes five paralogs unique to *S. hispanica*, which were duplicated after the divergence from *S. splendens*. The overactivity of this *SAD* gene tandem array produces an abundance of C18:1-ACP as a substrate for the desaturation of FAs in both the plastid and ER, and consequently the relatively high accumulation of ω-3 FAs in *S. hispanica* seeds. This contradicts the hypothesis that the high expression of ER localised *ShFAD3* drives the high ω-3 FA accumulation in *S. hispanica* seeds (Li et al., 2023). The WGD *ShSAD* genes (*ShSAD7*, *ShSAD11-a/b*, *ShSAD12*, and *ShSAD13*) have been under evolutionary constraints to maintain their function. On the other hand, *ShSAD* orthologs unique to *S. hispanica* (*ShSAD8*, *ShSAD9*, and *ShSAD10*) show diverging non-synonymous mutations and increased rate of substitution at specific sites (Kryazhimskiy & Plotkin, 2008) while showing evidence of purifying selection elsewhere.

## 5 CONCLUSIONS

This study generated a high-quality, chromosomal-level reference genome for *S. hispanica* to analyse the evolution of oil biosynthesis genes in this valuable oil-seed crop. Our analysis suggested that the expansion of the *ShSAD* gene family through tandem duplications is a driver of high ω-3 FAs accumulation in *S. hispanica* seeds. Comparative analysis of multiple chromosomal-level genomes is able to assess the putative effect of gene copy number variation and other source of structural genome variation. This work establishes valuable genomic resources in chia and prompts the need to investigate further structural variants at FA biosynthesis gene loci within and among species. This will enable the breeding of emerging crop or the horizontal transfer of genes across species, with the possibility of altering gene dosage through the introgression of arrays of paralogous genes to improve key traits.

## Supporting information

Shisp_Comparative_Genomics_Supplementary_Final

## ACKNOWLEDGMENTS

This research was supported by a Research Training Program Scholarship, the Alfred Nicholas Fellowship, the Megan Klemm Postgraduate Research Scholarship, and the Norma Hilda Schuster (nee Swift) Scholarship from the University of Melbourne awarded to TZ. BE was supported by the Inaugural Botany Foundation Fellowship 2020 from the University of Melbourne Botany Foundation.

## ORCID

**Tannaz Zare:** 0000-0002-7194-6800

**Jeff F. Paril:** 0000-0002-5693-4123

**Emma Barnett:** 0009-0002-6786-0972

**Parwinder Kaur:** 0000-0003-0201-0766

**Rudi Appels:** 0000-0002-5369-5227

**Berit Ebert:** 0000-0002-6914-5473

**Ute Roessner:** 0000-0002-6482-2615

**Alexandre Fournier-Level:** 0000-0002-6047-7164

## SUPPLEMENTAL MATERIAL

### Supplemental Figures

**Supplemental Figure S1:** Workflow of genomic DNA extraction from *S. hispanica*’s leaf tissue.

**Supplemental Figure S2:** An overview of the in situ Hi-C library preparation and sequencing.

**Supplemental Figure S3:** The Hi-C contact map of the *S. hispanica* genome assembly.

**Supplemental Figure S4:** Genome size estimation with GenomeScope using different k-mer lengths.

**Supplemental Figure S5:** Evaluation the completeness of the *S. hispanica* genome.

**Supplemental Figure S6:** Distribution of mapped tissue-specific transcripts from published assembled transcriptomes against the *S. hispanica* assembled genome.

**Supplemental Figure S7:** Distribution of repeat content and selected genomic features in each chromosome of *S. hispanica*.

**Supplemental Figure S8:** Gene Ontology (GO) enrichment analysis highlighting the top 10 enriched biological processes in the comparative genomics study of *S. hispanica* (p-value <0.05).

**Supplemental Figure S9**: Distribution of the Kernel density estimate (KDE) for transversion substitutions at fourfold degenerate sites (4dTV) for selected taxa studied here.

**Supplemental Figure S10:** Ka/Ks plots for three *ShSAD* genes of orthogroup OG0021142.

**Supplemental Figure S11:** Ka/Ks plots for three *ShSAD* genes of orthogroup OG0001529.

**Supplemental Figure S12:** Ka/Ks plots for a *ShSAD* gene (XP_047982897.1) and orthologous genes from *S. Splendens* belonging to orthogroup OG0001529.

**Supplemental Figure S13:** Ka/Ks plots for a ShSAD gene (XP_047982897.1) and orthologous genes from *A. thaliana* belonging to orthogroup OG0001529.

### Supplemental Tables

**Supplemental Table S1.** Percentage of trimmed paired-end Illumina reads with Trimmomatic

**Supplemental Table S2.** List of lipid genes and their functions used in whole genome comparative analysis of *S. hispanica*.

**Supplemental Table S3.** Summary statistics of inferred Hi-C contacts and mapped Hi-C reads to the *S. hispanica* draft assembly.

**Supplemental Table S4.** Summary statistics of the six pseudo-chromosomes of *S. hispanica* obtained by Hi-C scaffolding.

**Supplemental Table S5.** Identified proteins, RNA molecules, genes, and pseudogenes in *S. hispanica* chromosomes

**Supplemental Table S6.** Summary of repeat elements in the genome assembly of *S. hispanica*.

**Supplemental Table S7.** Gene Ontology (GO) enrichment analysis highlighting the top 20 enriched biological processes in the comparative genomics study of *S. hispanica* (p-value <0.05).

**Supplemental Table S8.** Gene Ontology (GO) enrichment analysis highlighting the top 20 enriched molecular functions in the comparative genomics study of *S. hispanica* (p-value <0.05).

**Supplemental Table S9.** Gene Ontology (GO) enrichment analysis highlighting the enriched cellular components in the comparative genomics study of *S. hispanica* (p-value <0.05).

## DATA AVAILABILITY

The Whole Genome Shotgun (WGS) project of *S. hispanica* is available at DDBJ/ENA/GenBank under the accession JALPBU000000000. The genome assembly and annotation of *S. hispanica* (NCBI *Salvia hispanica* Annotation Release 100) are available from the NCBI database under GenBank GCA_023119035.1 and RefSeq GCF_023119035.1 accessions.

