## Supplementary material for "Comparative Genomics Points to Tandem Duplications of *SAD* Gene Clusters as Drivers of Increased ω-3 Content in *S. hispanica* Seeds": Shisp_Comparative_Genomics_Supplementary_Final: Shisp_Comparative_Genomics_Supplementary_Final.pdf

Supplemental material consists of 21 pages, 13 supplemental figures and 9 supplemental tables.

### Supplemental Figures

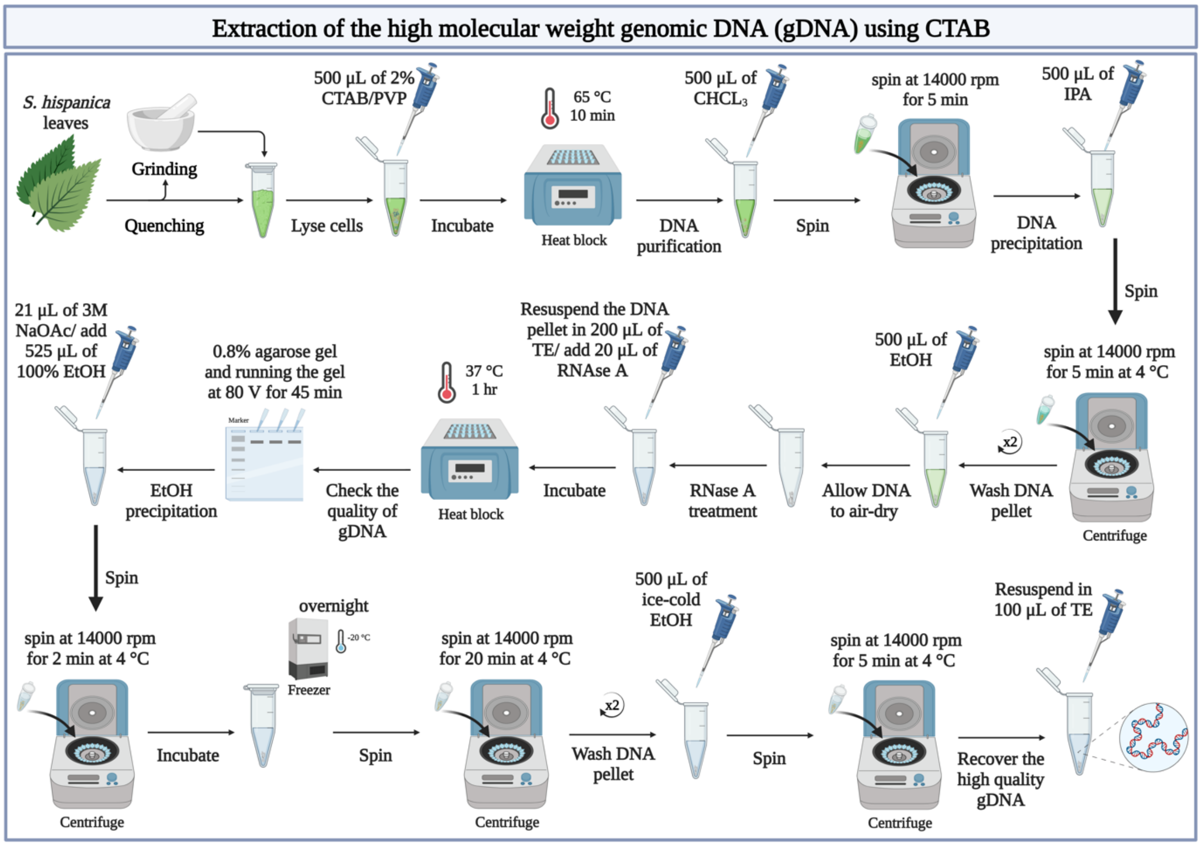


**Supplemental Figure S1.** Workflow of genomic DNA extraction from *S. hispanica*'s leaf tissue. The homogenised leaves were mixed with 500 µl of CTAB extraction buffer (2% w/v CTAB, 1% w/v polyvinylpyrrolidone (PVP-40mw), 20 mM ethylenediaminetetraacetic acid disodium salt (EDTA) pH 8.0, 100 mM Tris(hydroxymethyl)aminomethane (Tris-base) pH 8.0, and 1.5 M sodium chloride (NaCl)) and incubated for 10 min at 65 °C. The samples were mixed with 500 µl of chloroform (CHCl3) and centrifuged at 14,000 rpm for 5 min. The aqueous supernatant was transferred to a clean 1.5 ml Eppendorf tube, and an equal volume of isopropyl alcohol (C3H8O) was added, followed by gentle mixing for 2 min and centrifugation at 14,000 rpm for 5 min at 4 °C. The gDNA pellet was washed twice with 500 µl of 70% ethanol (EtOH) followed by centrifugation at 14,000 rpm for 5 min at 4 °C. The EtOH was discarded, and the tube was left open overnight to evaporate the residual EtOH. The dried gDNA pellet was resuspended in 100 µl of Tris-EDTA buffer (pH 8.0). Created with BioRender.com.

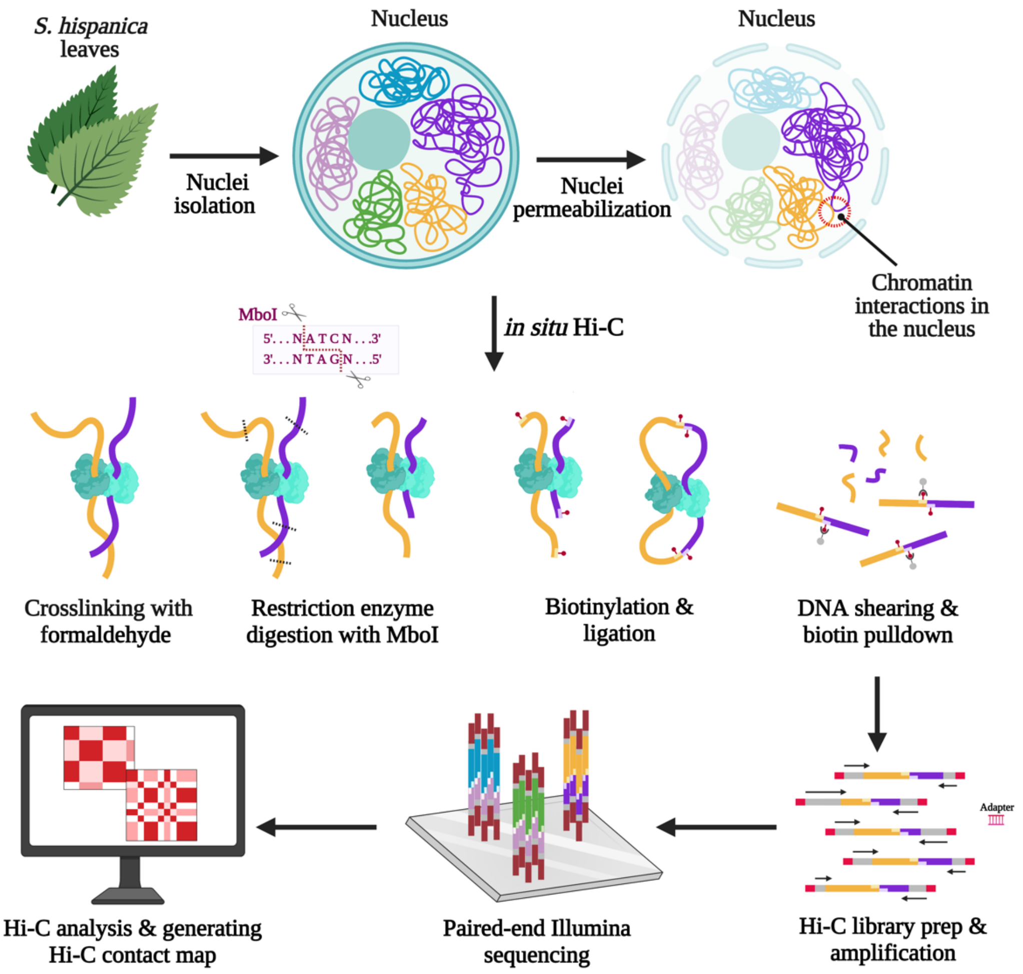


**Supplemental Figure S2.** An overview of the *in situ* Hi-C library preparation and sequencing. Green and healthy-looking leaves of *S. hispanica* were fixed with 1% formaldehyde solution at RT. Fixed tissues were resuspended in a nuclei isolation buffer and homogenised with liquid nitrogen. Following nuclei isolation, the DNA-DNA proximity ligation is performed in intact nuclei using formaldehyde (i.e., nuclei permeabilisation & formaldehyde-crosslinking). The crosslinked chromatin fragments were then digested by a four-cutter restriction enzyme (*MboI*, with GATCGATC ligation junction sequence), followed by biotinylation to label the 5' and 3' restriction fragment overhangs. The biotinylated fragments were then ligated, and crosslinks were reversed to remove protein contaminations and unligated fragments. The resulting ligated DNA fragments were purified and sheared to allow for the biotin pull down of the point ligation junctions using streptavidin beads prior to the sequencing. The index adapters were added to the fragments, and the Hi-C library was amplified with PCR using Illumina primers for paired-end Illumina sequencing. The sequencing data were collected and processed through the Juicer and 3D-DNA pipelines to construct the Hi-C contact maps for the *S. hispanica* genome. The experimental protocol was adopted from Rao et al. (2014). The figure was created with BioRender.com.

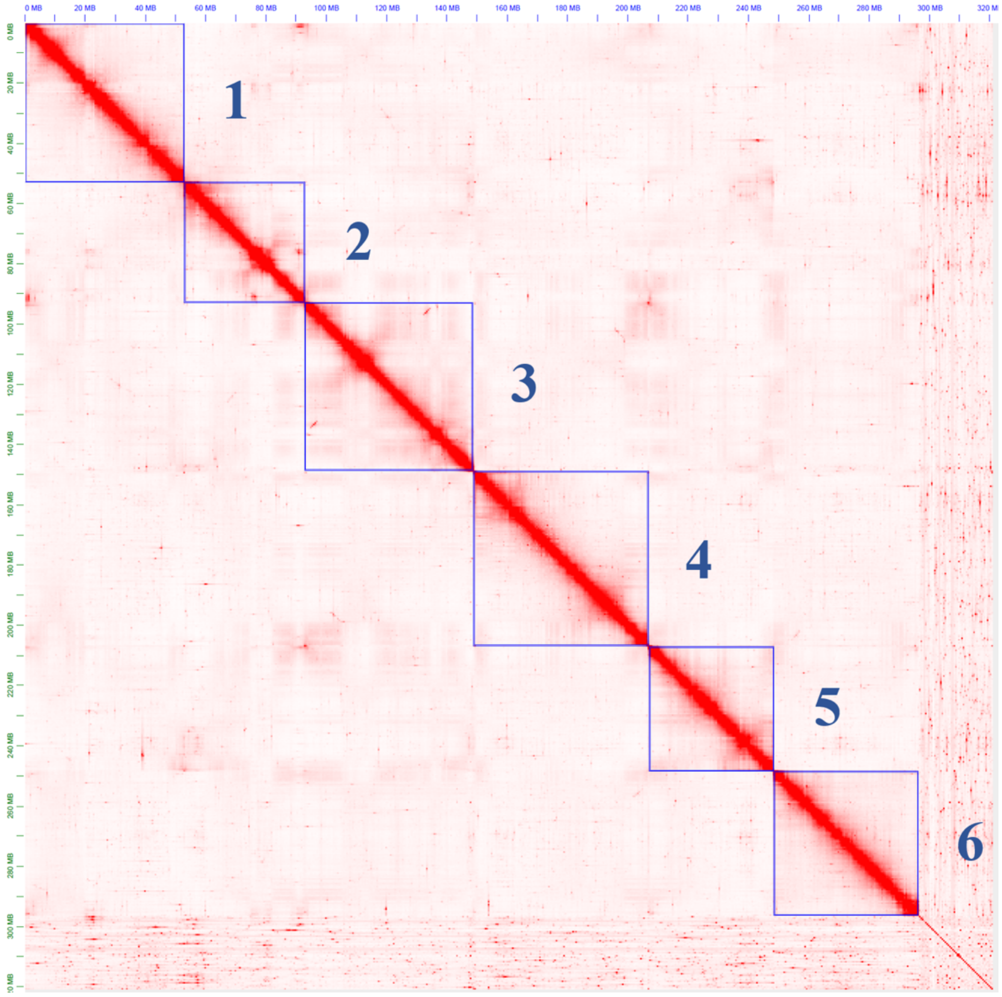


**Supplemental Figure S3.** The Hi-C contact map of the *S. hispanica* genome assembly. The contact map was visualised using Juicebox (v1.11.08) at the resolution of 500 kbp with balanced normalisation and colour range from 0 to 5000 for the observed counts. The six pseudo-chromosomes of *S. hispanica* are enclosed in blue boxes.

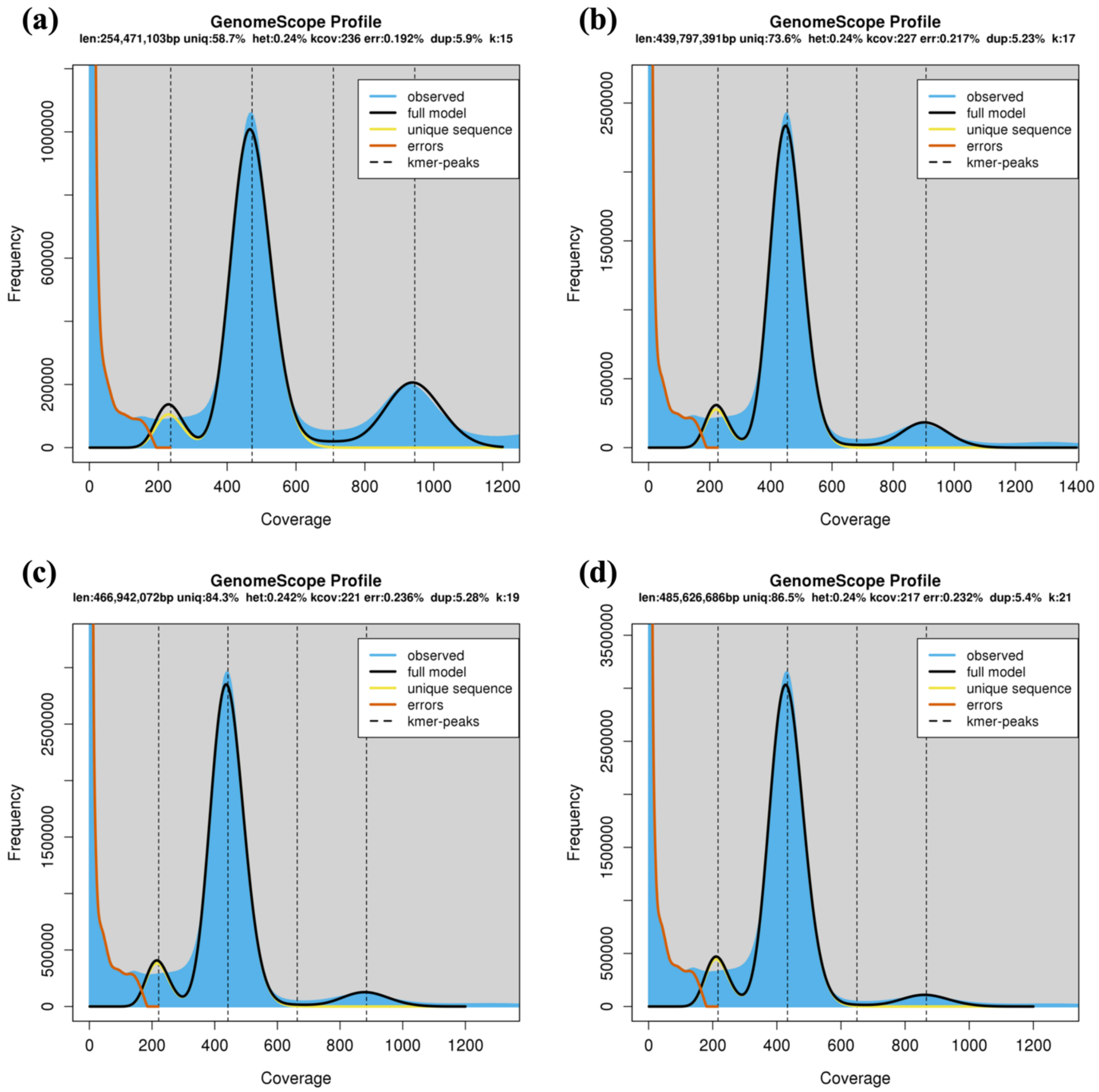


**Supplemental Figure S4.** Genome size estimation with GenomeScope using different k-mer lengths. GenomeScope calculates the k-mer frequency distribution of (a) 15-mer, (b) 17-mer, (c) 19-mer, and (d) 21-mer across the Illumina reads. Haploid genome length (len); length of unique sequences (uniq); estimated heterozygosity (het); mean k-mer coverage (kcov); error rate of sequencing reads (err); duplication rate (dup); k-mer length (k).

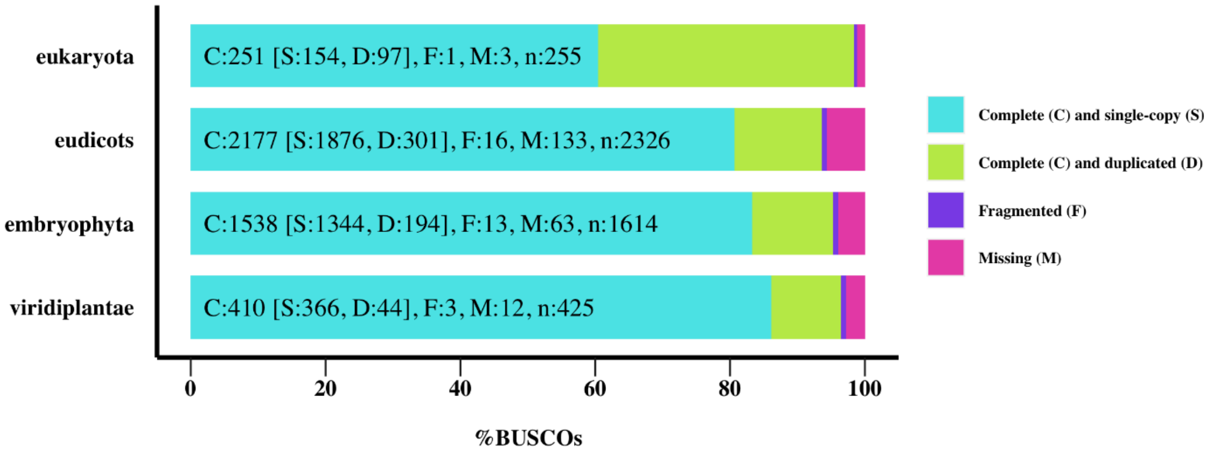


**Supplemental Figure S5.** Evaluation of the completeness of the *S. hispanica* genome. (A) BUSCO. analysis using different lineages; Complete (C) and single-copy (S); Duplicated (D); Fragmented (F); Missing (M); Total number of genes (n).

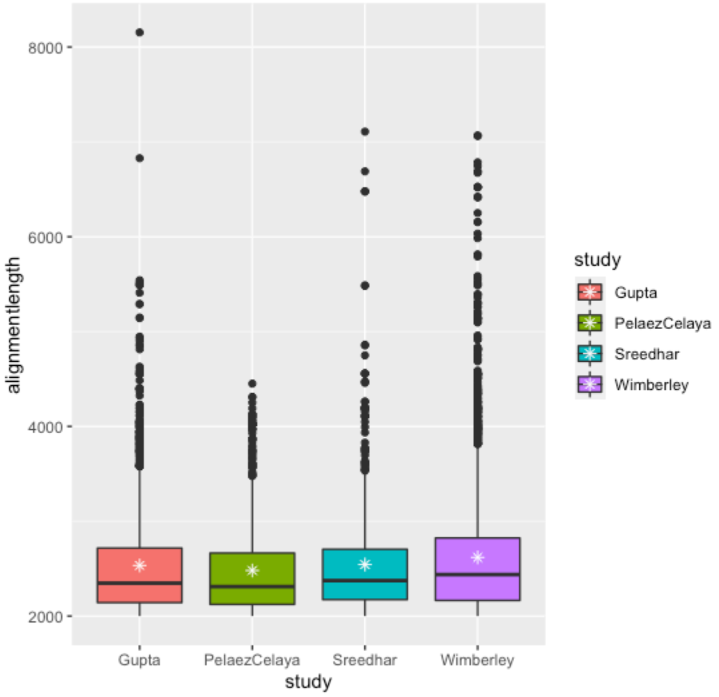


**Supplemental Figure S6.** Distribution of mapped tissue-specific transcripts from published assembled transcriptomes against the *S. hispanica* assembled genome. Box plots indicate the highest mean and median alignment length when comparing the Wimberley study transcripts with the assembled genome.

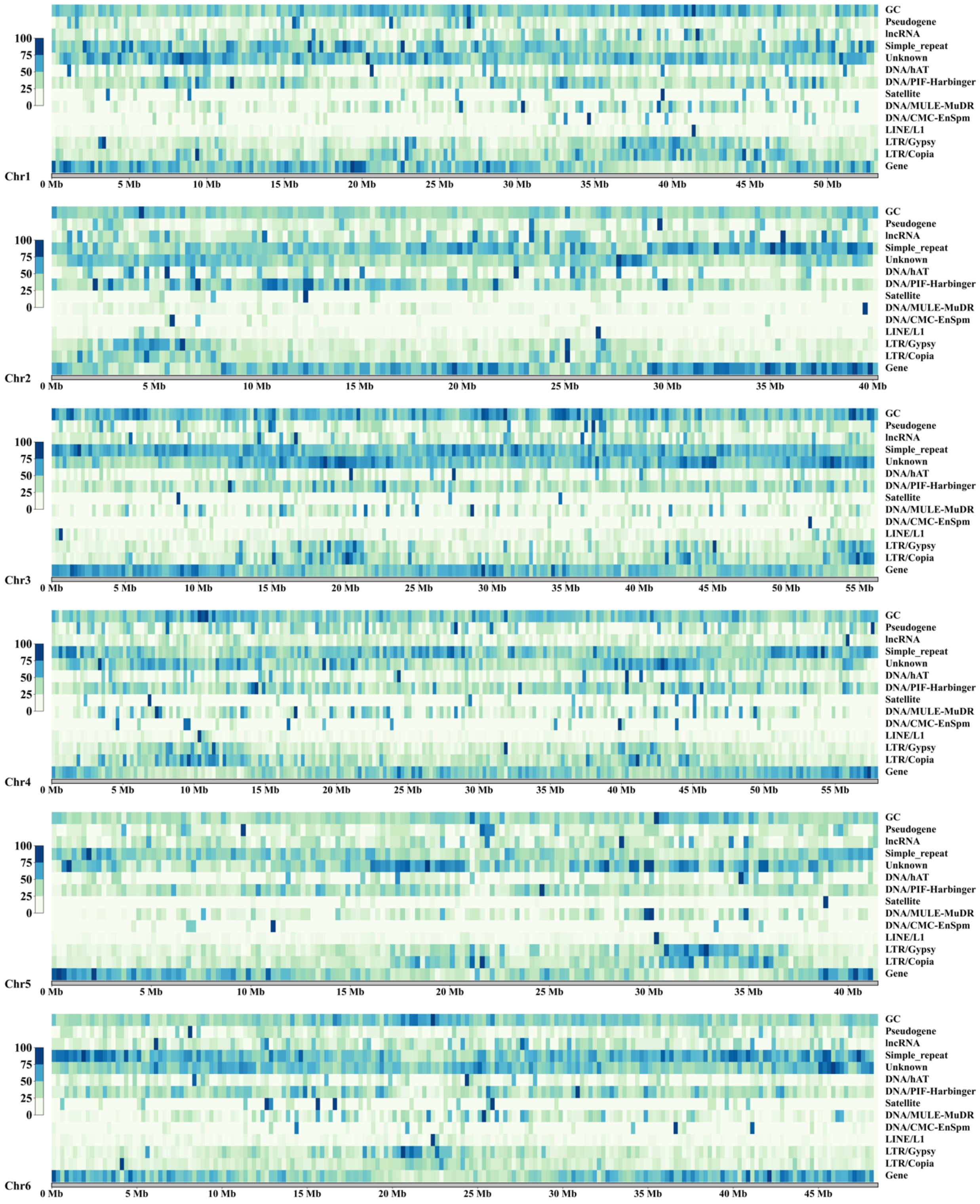


**Supplemental Figure S7.** Distribution of repeat content and selected genomic features in each chromosome of *S. hispanica* are reported as percentages at a window size of 250 kbp and shown as density heatmaps. Gene refers to protein coding genes. The position of centromeric and telomeric regions within each chromosome can be inferred from the intensity of features. Centromeric regions show higher GC%, Gypsy and Copia intensity, while telomeric regions show the higher intensity of gene and simple repeat content.

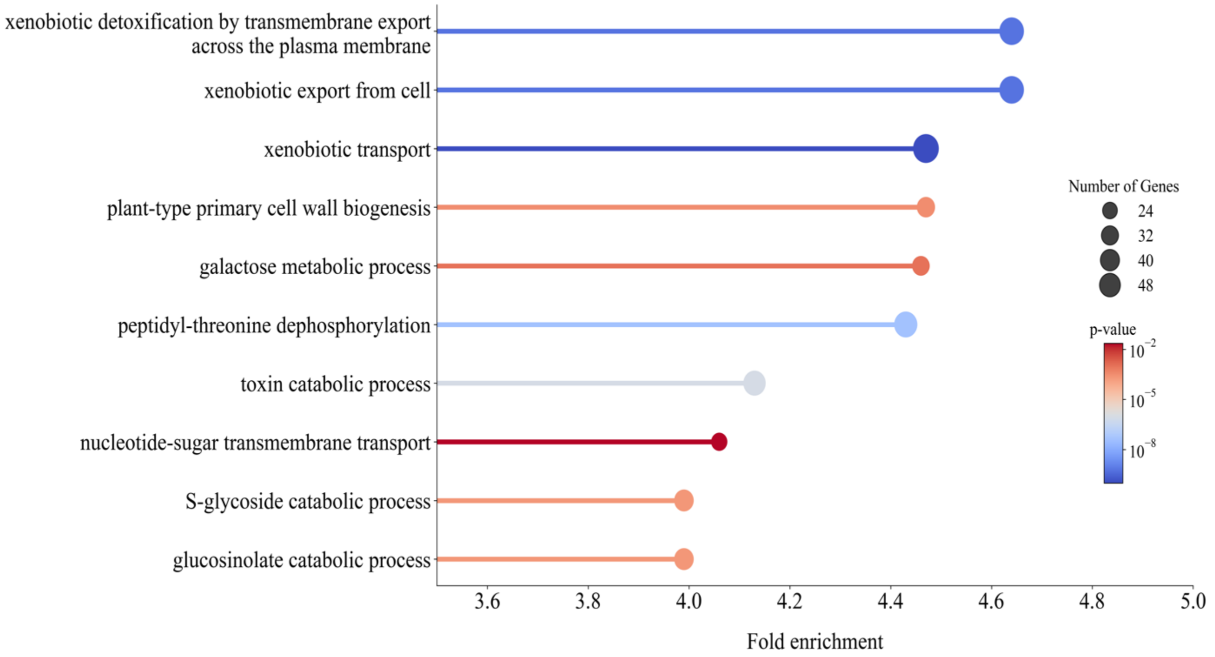


**Supplemental Figure S8.** Gene Ontology (GO) enrichment analysis highlighting the top 10 enriched biological processes in the comparative genomics study of *S. hispanica* (p-value <0.05).

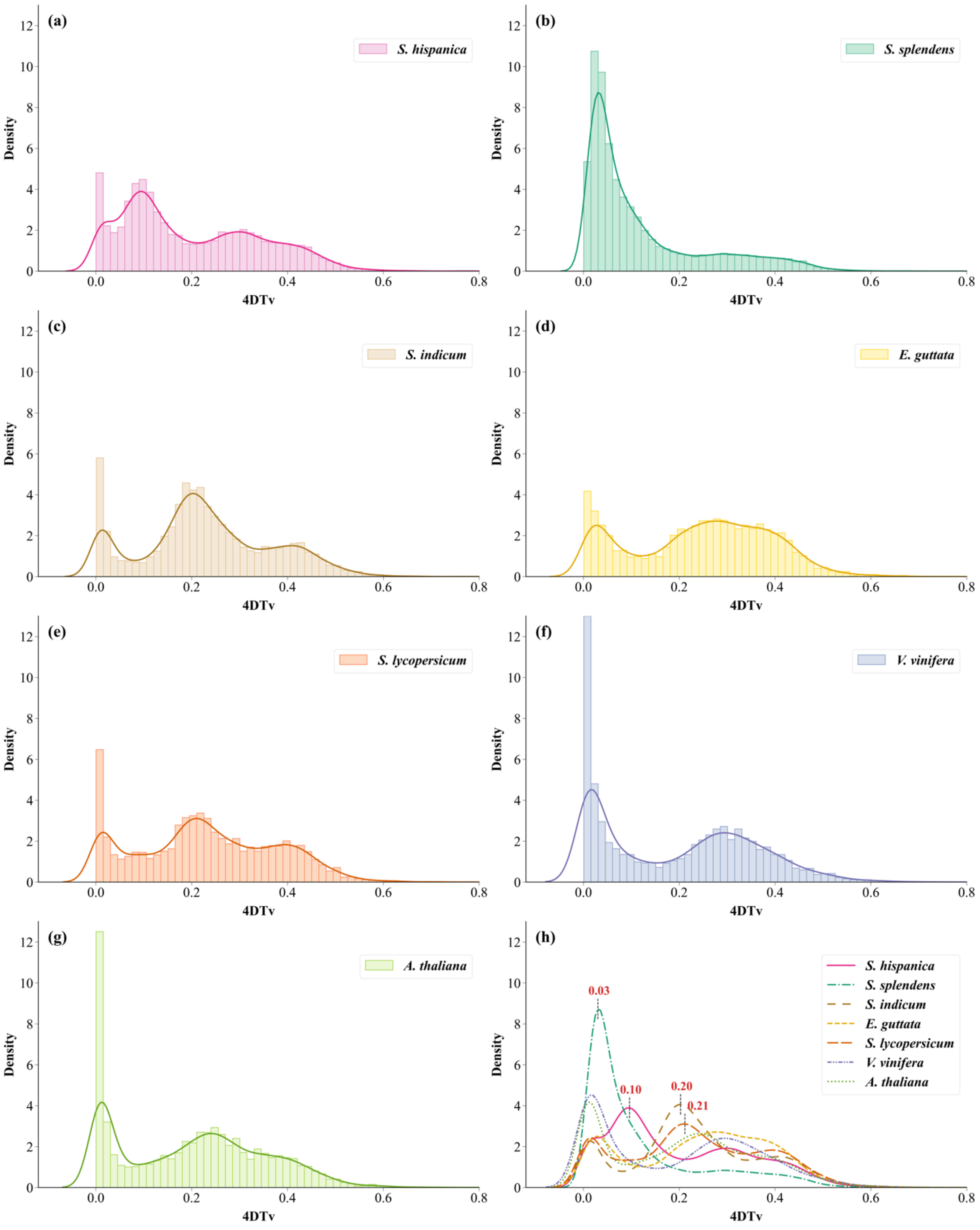


**Supplemental Figure S9.** Distribution of the Kernel density estimate (KDE) for transversion substitutions at fourfold degenerate sites (4dTV) for selected taxa studied here. (a-g) The curves of the kernel density estimate (KDE) overlapped with the distribution of 4DTv for each species. (h) Comparison of the KDE of all species. Vertical lines indicate WGD events.

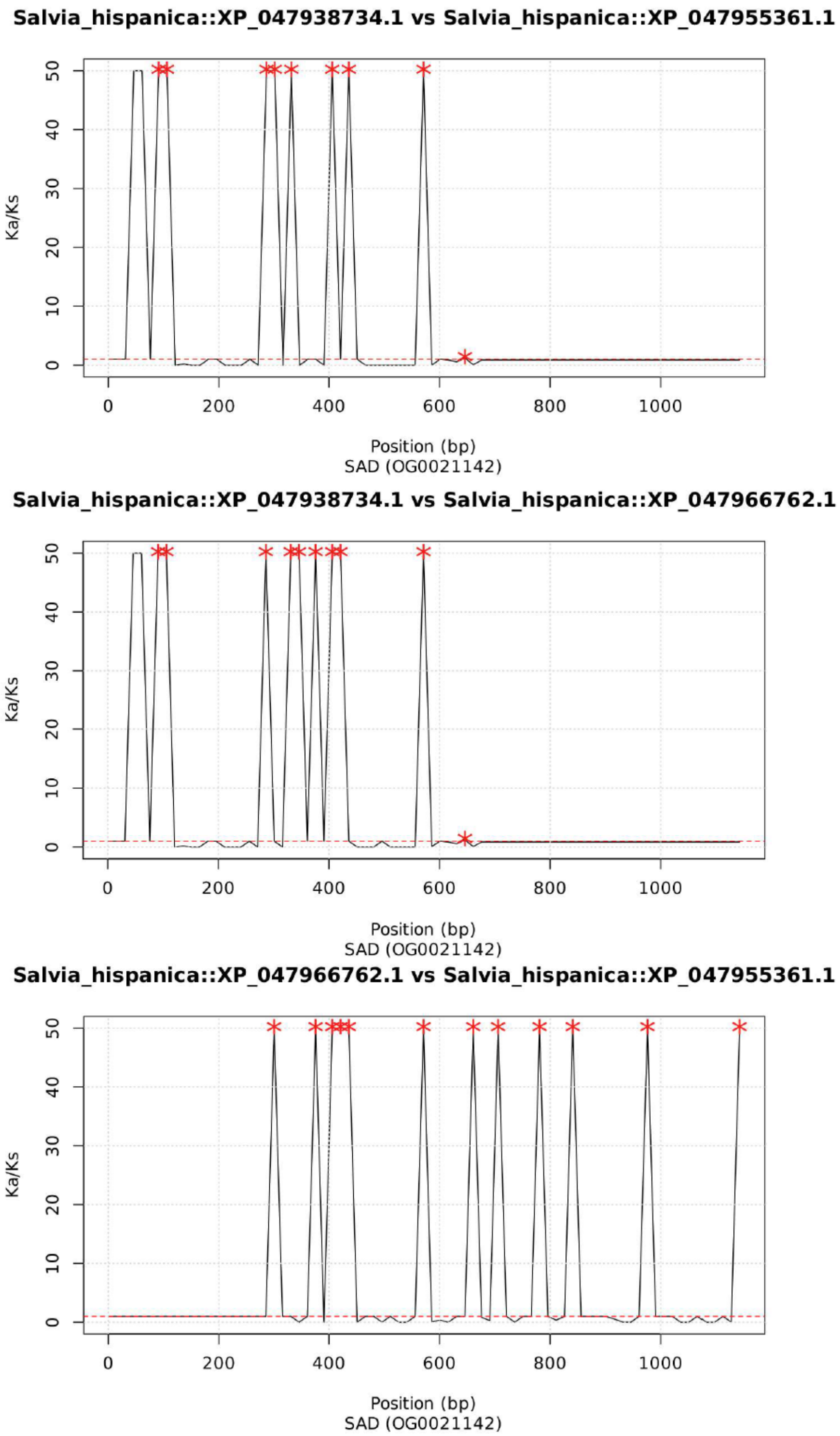


**Supplemental Figure S10.** Ka/Ks plots for three *ShSAD* genes of orthogroup OG0021142. XP identifiers refer to the *S. hispanica* proteins. Stars indicate statistically significant peaks. The horizontal red dashed line indicates the Ka/Ks ratio of 1. XP_047966762.1: *ShSAD8*; XP_047955361.1: *ShSAD9*; XP_047938734.1: *ShSAD10*.

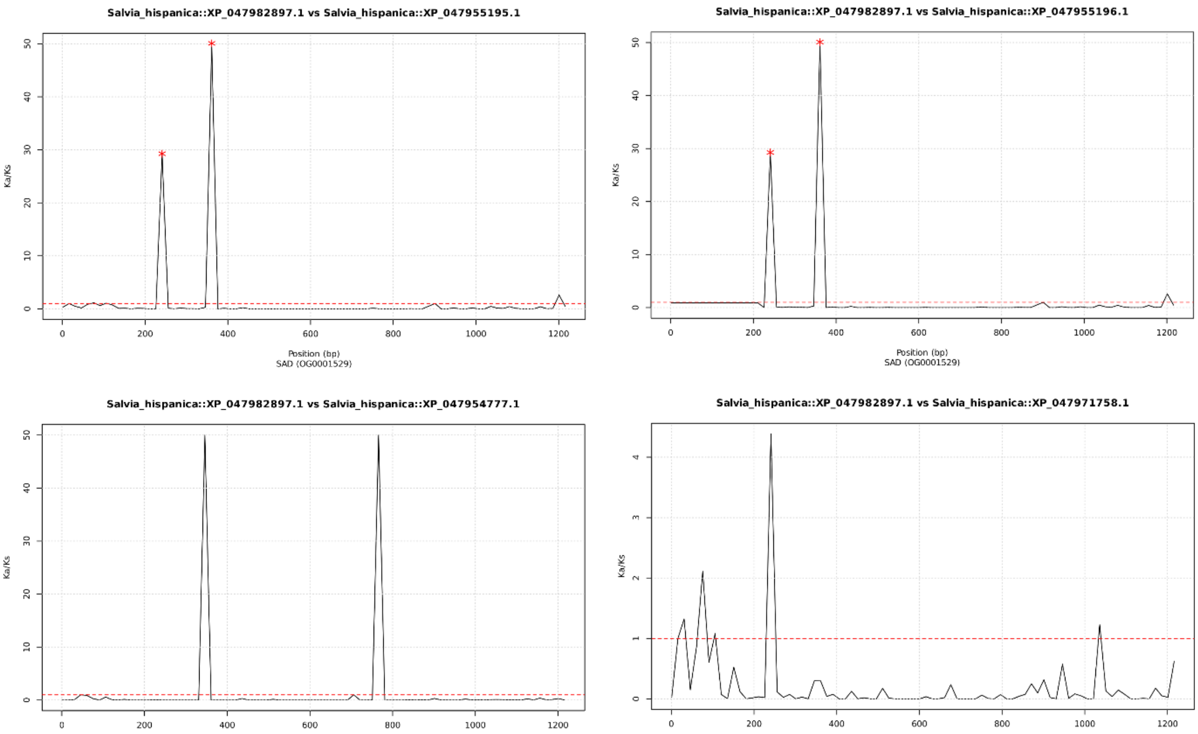


**Supplemental Figure S11.** Ka/Ks plots for three *ShSAD* genes of orthogroup OG0001529. XP identifiers refer to the *S. hispanica* proteins. Stars indicate statistically significant peaks. The horizontal red dashed line indicates the Ka/Ks ratio of 1. XP_047982897.1: *ShSAD13*; XP_047955195.1: *ShSAD11-a* isoform X1; XP_047955596.1: *ShSAD2*; XP_047954777.1: *ShSAD7*; XP_047971758.1: *ShSAD12*.

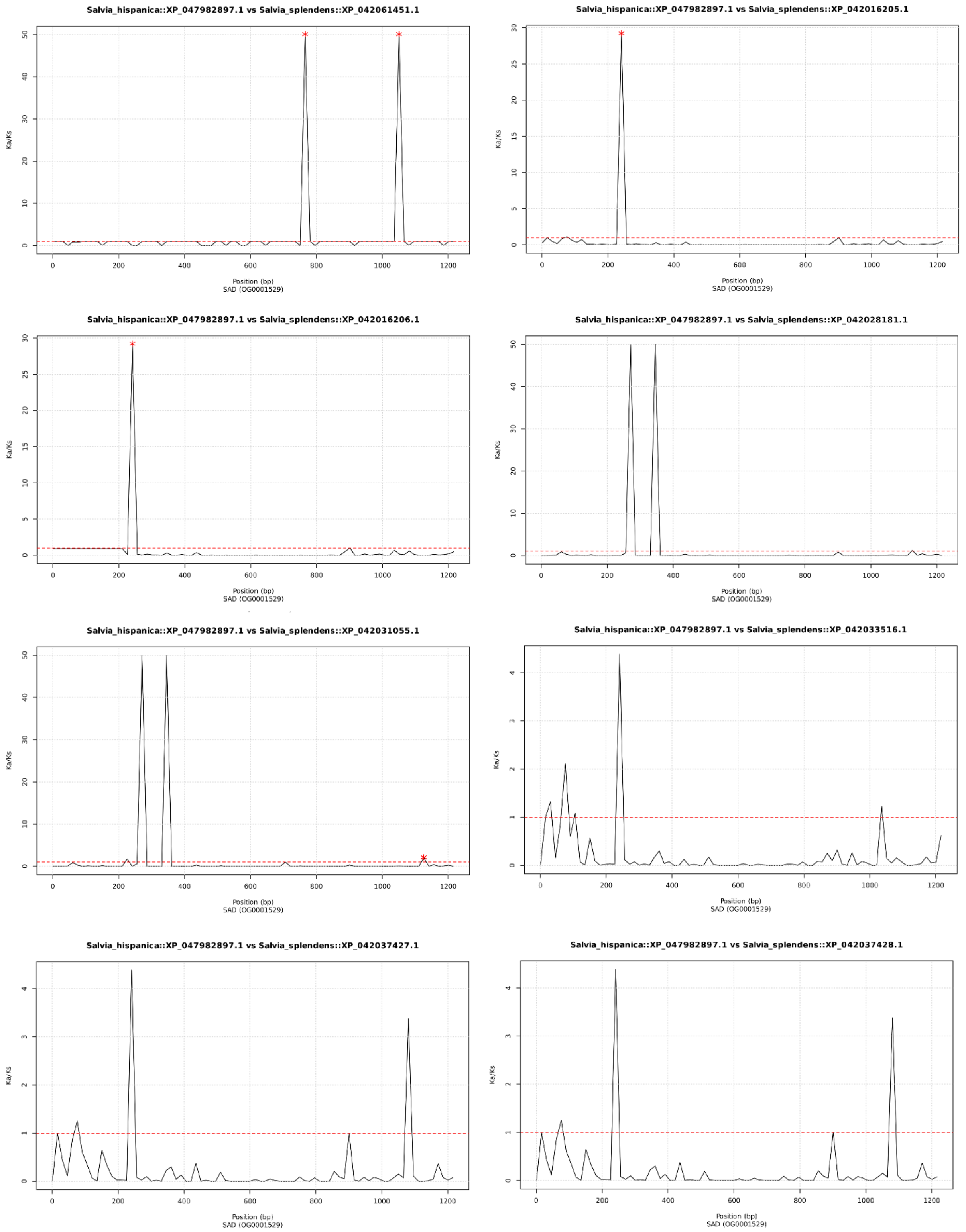


**Supplemental Figure S12.** Ka/Ks plots for a *ShSAD* gene (XP_047982897.1) and orthologous genes from *S. Splendens* belonging to orthogroup OG0001529. XP identifiers refer to the *S. hispanica* proteins. Stars indicate statistically significant peaks. The horizontal red dashed line indicates a Ka/Ks ratio of 1. XP_047982897.1: *ShSAD13*.

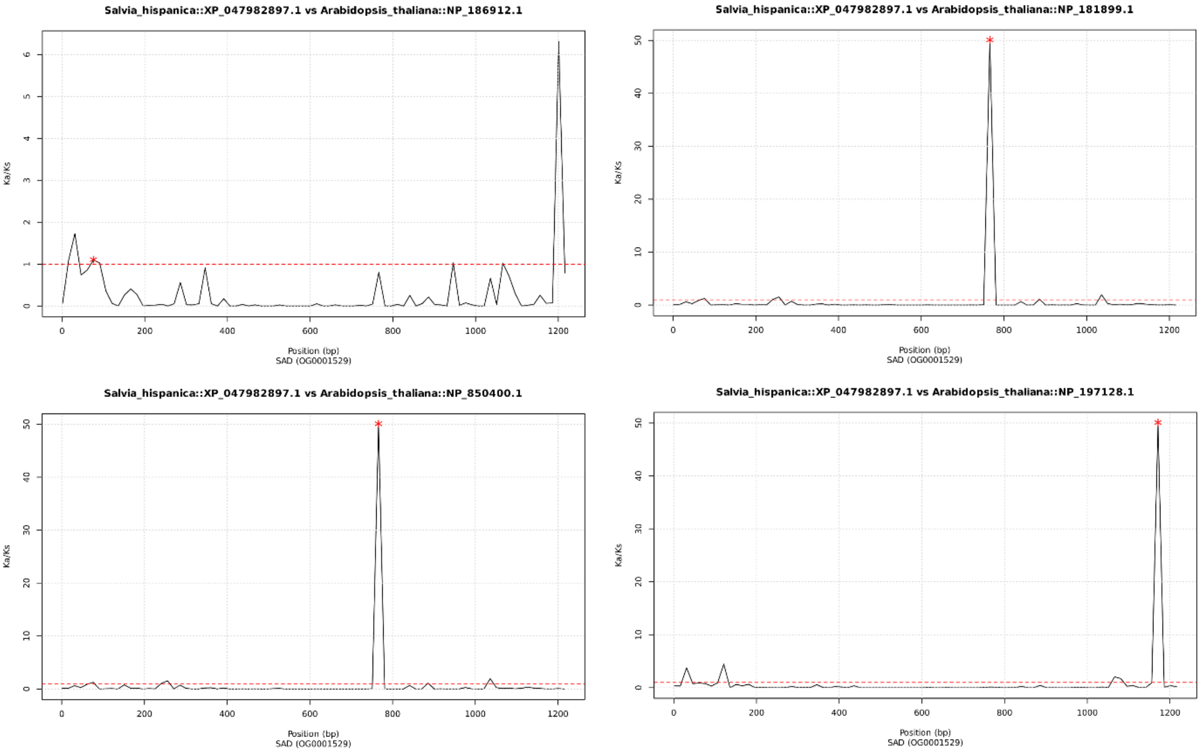


**Supplemental Figure S13.** Ka/Ks plots for a *ShSAD* gene (XP_047982897.1) and orthologous genes from *A. thaliana* belonging to orthogroup OG0001529. XP identifiers refer to the *S. hispanica* proteins. Stars indicate peaks that are statistically significant. The horizontal red dashed line indicates a Ka/Ks ratio of 1. XP_047982897.1: *ShSAD13*.

### Supplemental Tables

**Supplemental Table S1.** Percentage of trimmed paired-end Illumina reads with Trimmomatic

| Paired-end libraries | Input read pairs (raw) | Surviving reads | Forward only survived reads | Reverse only survived reads | Discarded reads |
|---|---|---|---|---|---|
| **s1_combined_trimmed_R1.fastq** | 1677484634 | 1476561377 | 119666880 | 40166588 | 41089789 |
| **s1_combined_trimmed_R2.fastq** | | (88.02%) | (7.13%) | (2.39%) | (2.45%) |

**Supplemental Table S2.** List of lipid genes and their functions used in whole genome comparative analysis of *S. hispanica*.

| Gene Name | Gene Symbol |
|---|---|
| Acetyl-CoA carboxylase | *ACC* |
| Acyl carrier protein | *ACP* |
| Biotin carboxylase | *BC* |
| Biotin carboxyl carrier protein of acetyl (CoA) carboxylase 1 | *BCCP1* |
| Biotin carboxyl carrier protein of acetyl (CoA) carboxylase 2 | *BCCP2* |
| acetyl-CoA carboxyltransferase subunit α | *α-CT* |
| acetyl-CoA carboxyltransferase subunit β | *β-CT* |
| Diacylglycerol O-acyltransferase | *DGAT* |
| Enoyl-acyl carrier protein (enoyl-ACP) reductase | *ENR* |
| Microsomal ω-6 fatty acid desaturase 2 | *FAD2* |
| Microsomal ω-3 fatty acid desaturase 3 | *FAD3* |
| Plastidial fatty acid desaturase 4 | *FAD4* |
| Plastidial Palmitoyl-monogalactosyldiacylglycerol Delta7-desaturase | *Fad5* |
| Plastidial ω-6 fatty acid desaturase 6 | *FAD6* |
| Plastidial ω-3 fatty acid desaturase 7 | *FAD7* |
| Temperature-sensitive plastidial ω -3 fatty acid desaturase 8 | *FAD8* |
| Fatty acyl- acyl carrier protein (ACP) thioesterase A | *FATA* |
| Fatty acyl- acyl carrier protein (ACP) thioesterase B | *FATB* |
| Glycerol-3-phosphate acyltransferase | *GPAT* |
| 3-hydroxyacyl- acyl carrier protein (ACP) dehydratase | *HAD* |
| 3-oxoacyl- acyl carrier protein (ACP) reductase | *KAR* |
| β-ketoacyl- acyl carrier protein (ACP) synthase I | *KAS I* |
| β-ketoacyl- acyl carrier protein (ACP) synthase II | *KAS II* |
| β-ketoacyl- acyl carrier protein (ACP) synthase III | *KAS III* |
| Long-chain acyl-coenzyme A synthetase | *LACS* |
| Lysophosphatidic acid acyltransferase | *LPAAT* |
| Lysophosphatidylcholine acyltransferase | *LPCAT* |
| Non-specific lipid-transfer type 1 | *LPT1* |
| Lipoxygenase | *LOX* |
| Malonyl-CoA: acyl carrier protein (ACP) transacylase | *MCAT* |
| Phosphatidic acid phosphatase | *PAP* |
| Phospholipid:diacylglycerol acyltransferase | *PDAT* |
| Phosphatidylcholine:diacylglycerol cholinephosphotransferases | *PDCT* |
| Phosphatidylglycerophosphate synthase | *PGPS* |
| Stearoyl-acyl carrier protein desaturase | *SAD* |

**Supplemental Table S3.** Summary statistics of inferred Hi-C contacts and mapped Hi-C reads to the *S. hispanica* draft assembly.

| Hi-C statistics type | Statistics |
|---|---|
| Sequenced Read Pairs | 1,425,815,599 |
| Normal Paired | 580,212,682 (40.69%) |
| Chimeric Paired | 523,807,128 (36.74%) |
| Chimeric Ambiguous | 281,548,738 (19.75%) |
| Unmapped | 40,247,051 (2.82%) |
| Ligation Motif Present | 0 (0.00%) |
| Alignable (Normal + Chimeric Paired) | 1,104,019,810 (77.43%) |
| Unique Reads | 378,602,733 (26.55%) |
| PCR Duplicates | 722,921,771 (50.70%) |
| Optical Duplicates | 2,495,306 (0.18%) |
| Library Complexity Estimate | 405,380,268 |
| Intra-fragment Reads | 25,592,418 (1.79% / 6.76%) |
| Below MAPQ Threshold | 179,140,938 (12.56% / 47.32%) |
| Hi-C Contacts | 173,869,377 (12.19% / 45.92%) |
| Ligation Motif Present | 0 (0.00% / 0.00%) |
| 3' Bias (Long Range) | 49% - 51% |
| Pair Type %(L-I-O-R) | 25% - 25% - 25% - 25% |
| Inter-chromosomal | 127,912,572 (8.97% / 33.79%) |
| Intra-chromosomal | 45,956,805 (3.22% / 12.14%) |
| Short Range (<20Kb) | 39,258,468 (2.75% / 10.37%) |
| Long Range (>20Kb) | 6,698,144 (0.47% / 1.77%) |

**Supplemental Table S4.** Summary statistics of the six pseudo-chromosomes of *S. hispanica* obtained by Hi-C scaffolding.

| Chromosome | Total Length (Mb) | Ungapped Length (Mb) | N50 | Spanned Gap (bp) | GC% |
|---|---|---|---|---|---|
| Chr 1 | 53.19 | 53.06 | 53.19 | 697 | 35.3 |
| Chr 2 | 40.18 | 40.08 | 40.18 | 536 | 36.1 |
| Chr 3 | 56.04 | 55.97 | 56.04 | 558 | 35.8 |
| Chr 4 | 57.98 | 57.87 | 57.98 | 741 | 36.1 |
| Chr 5 | 41.45 | 41.38 | 41.45 | 491 | 36.4 |
| Chr 6 | 48.31 | 48.21 | 48.31 | 703 | 36.4 |

**Supplemental Table S5.** Identified proteins, RNA molecules, genes, and pseudogenes in *S. hispanica* chromosomes

| Chromosome | Protein | rRNA | tRNA | Other RNA | Gene | Pseudogene |
|---|---|---|---|---|---|---|
| Chr 1 | 7,239 | - | 109 | 1,121 | 6,205 | 359 |
| Chr 2 | 6,287 | - | 96 | 884 | 5,172 | 254 |
| Chr 3 | 9,601 | 1 | 179 | 1,266 | 7,770 | 365 |
| Chr 4 | 8,676 | - | 126 | 1,284 | 7,207 | 370 |
| Chr 5 | 6,181 | 7 | 75 | 821 | 5,022 | 226 |
| Chr 6 | 6,842 | 24 | 82 | 988 | 5641 | 270 |

**Supplemental Table S6.** Summary of repeat elements in the genome assembly of *S. hispanica*.

| Family | Element | Number of elements | Length occupied | Percent of Genome |
|---|---|---|---|---|
| **DNA** | CMC-EnSpm | 374 | 173,908 (bp) | 0.05 % |
| | MULE-MuDR | 3,765 | 2,053,776 (bp) | 0.64 % |
| | PIF-Harbinger | 15,734 | 8,207,998 (bp) | 2.55 % |
| | TcMar-Stowaway | 582 | 247,182 (bp) | 0.08 % |
| | TcMar-Tc4 | 1,442 | 240,647 (bp) | 0.07 % |
| | hAT | 391 | 54,264 (bp) | 0.02 % |
| | hAT-Ac | 1,641 | 599,301 (bp) | 0.19 % |
| | hAT-Tip100 | 817 | 284,702 (bp) | 0.09 % |
| **LINE** | L1 | 1,981 | 1,227,606 (bp) | 0.38 % |
| **LTR** | Copia | 20,837 | 17,993,947 (bp) | 5.60% |
| | Gypsy | 18,472 | 17,519,657 (bp) | 5.45 % |
| | LTR/Unknown | 29,744 | 7,997,371 (bp) | 2.49 % |
| **SINE** | | 63 | 4,537 (bp) | 0.00 % |
| **Rolling circles (RC)** | Helitron | 3,894 | 2,032,369 (bp) | 0.63 % |
| **Satellites** | | 316 | 64,157 (bp) | 0.02 % |
| **Simple repeats** | | 61,732 | 2,569,321 (bp) | 0.80 % |
| **Low complexity** | | 12,704 | 603,934 (bp) | 0.19 % |
| **rRNA** | | 850 | 131,804 (bp) | 0.04 % |
| **tRNA** | | 906 | 131,766 (bp) | 0.04 % |
| **snRNA** | | 400 | 70,348 (bp) | 0.02 % |
| **Unclassified** | | 349,108 | 79,678,338 (bp) | 24.79 % |
| **Total repeats** | | | 141,886,933 (bp) | 44.14 % |

**Supplemental Table S7.** Gene Ontology (GO) enrichment analysis highlighting the top 20 enriched biological processes in the comparative genomics study of *S. hispanica* (p-value <0.05).

| GO ID | GO Biological Process | Fold enrichment | p-value Bonferroni |
|---|---|---|---|
| GO:1990961 | xenobiotic detoxification by transmembrane export | 4.64 | 5.70E-10 |
| GO:0046618 | xenobiotic export from cell | 4.64 | 5.70E-10 |
| GO:0042908 | xenobiotic transport | 4.47 | 9.63E-11 |
| GO:0009833 | plant-type primary cell wall biogenesis | 4.47 | 3.22E-04 |
| GO:0006012 | galactose metabolic process | 4.46 | 9.83E-04 |
| GO:0035970 | peptidyl-threonine dephosphorylation | 4.43 | 4.17E-08 |
| GO:0009407 | toxin catabolic process | 4.13 | 8.70E-07 |
| GO:0015780 | nucleotide-sugar transmembrane transport | 4.06 | 2.30E-02 |
| GO:0016145 | S-glycoside catabolic process | 3.99 | 2.14E-04 |
| GO:0019762 | glucosinolate catabolic process | 3.99 | 2.14E-04 |
| GO:0140115 | export across plasma membrane | 3.71 | 5.33E-11 |
| GO:0006885 | regulation of pH | 3.48 | 9.27E-06 |
| GO:0044772 | mitotic cell cycle phase transition | 3.48 | 6.59E-05 |
| GO:1901264 | carbohydrate derivative transport | 3.44 | 2.43E-05 |
| GO:0044770 | cell cycle phase transition | 3.35 | 1.74E-04 |
| GO:0000281 | mitotic cytokinesis | 3.31 | 7.88E-03 |
| GO:0007166 | cell surface receptor signalling pathway | 3.27 | 5.13E-05 |
| GO:0003333 | amino acid transmembrane transport | 3.25 | 1.78E-04 |
| GO:0098754 | detoxification | 3.19 | 3.68E-15 |
| GO:1903825 | organic acid transmembrane transport | 3.15 | 1.59E-06 |

**Supplemental Table S8.** Gene Ontology (GO) enrichment analysis highlighting the top 20 enriched molecular functions in the comparative genomics study of *S. hispanica* (p-value <0.05).

| GO ID | GO Molecular Functions | Fold enrichment | p-value Bonferroni |
|---|---|---|---|
| GO:0004033 | aldo-keto reductase (NADP) activity | 4.64 | 1.90E-03 |
| GO:0000293 | ferric-chelate reductase activity | 4.64 | 1.03E-02 |
| GO:0030570 | pectate lyase activity | 4.64 | 3.53E-04 |
| GO:0042910 | xenobiotic transmembrane transporter activity | 4.57 | 2.95E-15 |
| GO:0016760 | cellulose synthase (UDP-forming) activity | 4.46 | 5.76E-04 |
| GO:0102483 | scopolin beta-glucosidase activity | 4.33 | 1.18E-07 |
| GO:0005200 | structural constituent of cytoskeleton | 4.29 | 9.23E-04 |
| GO:0005338 | nucleotide-sugar transmembrane transporter activity | 4.1 | 4.43E-03 |
| GO:0080044 | quercetin 7-O-glucosyltransferase activity | 4.05 | 2.63E-08 |
| GO:0004707 | MAP kinase activity | 4.03 | 2.36E-02 |
| GO:0008810 | cellulase activity | 4 | 2.25E-03 |
| GO:0015165 | pyrimidine nucleotide-sugar transmembrane transporter | 4 | 4.13E-02 |
| GO:0016413 | O-acetyltransferase activity | 3.94 | 2.08E-07 |
| GO:0000156 | phosphorelay response regulator activity | 3.93 | 6.56E-04 |
| GO:1903231 | mRNA base-pairing post-transcriptional repressor | 3.92 | 1.19E-02 |
| GO:0051119 | sugar transmembrane transporter activity | 3.89 | 1.06E-08 |
| GO:0016538 | cyclin-dependent protein serine/threonine kinase regulator | 3.83 | 9.41E-07 |
| GO:0016857 | racemase and epimerase activity | 3.82 | 9.94E-04 |
| GO:0042562 | hormone binding | 3.79 | 1.72E-03 |
| GO:0016722 | oxidoreductase activity, acting on metal ions | 3.78 | 1.80E-02 |

**Supplemental Table S9.** Gene Ontology (GO) enrichment analysis highlighting the enriched cellular components in the comparative genomics study of *S. hispanica* (p-value <0.05).

| GO ID | GO Cellular Components | Fold enrichment | p-value Bonferroni |
|---|---|---|---|
| GO:0005764 | lysosome | 3.42 | 5.11E-04 |
| GO:0000323 | lytic vacuole | 2.99 | 6.22E-03 |
| GO:0031985 | Golgi cisterna | 1.85 | 1.03E-02 |
| GO:0005795 | Golgi stack | 1.81 | 1.52E-02 |
| GO:0005794 | Golgi apparatus | 1.53 | 3.11E-10 |
| GO:0016021 | integral component of membrane | 1.46 | 1.57E-02 |
| GO:0005886 | plasma membrane | 1.36 | 1.49E-14 |
| GO:0071944 | cell periphery | 1.3 | 8.46E-12 |
| GO:0012505 | endomembrane system | 1.27 | 1.25E-05 |
| GO:0016020 | membrane | 1.26 | 1.39E-13 |
| GO:0009507 | chloroplast | 1.16 | 7.97E-05 |
| GO:0009536 | plastid | 1.15 | 4.15E-04 |
| GO:0005737 | cytoplasm | 1.1 | 2.08E-11 |
| GO:0110165 | cellular anatomical entity | 1.04 | 6.83E-16 |
| GO:0005622 | intracellular anatomical structure | 1.03 | 1.42E-03 |
